# Specific photoreceptor cell fate pathways are differentially altered in NR2E3-associated diseases

**DOI:** 10.1101/2023.06.03.543550

**Authors:** Izarbe Aísa-Marín, Quirze Rovira, Noelia Díaz, Laura Calvo-López, Juan M. Vaquerizas, Gemma Marfany

**Affiliations:** Department de Genètica, Microbiologia i Estadística, Universitat de Barcelona, Barcelona, 08028, Spain; IBUB-IRSJD, Institut de Biomedicina de la Universitat de Barcelona-Institut de Recerca Sant Joan de Déu, Barcelona, 08028, Spain; CIBERER, Instituto de Salud Carlos III, Barcelona, 08028, Spain; Max-Planck-Institute for Molecular Biomedicine, Münster, 48149, Germany; Current position: Institute of Marine Sciences, The Spanish National Research Council (CSIC), 08003, Spain; MRC London Institute of Medical Sciences, Institute of Clinical Sciences, Imperial College London, London, W12 0NN, UK; Institute of Clinical Sciences, Faculty of Medicine, Imperial College London, Du Cane Road, London W12 0NN, UK; DBGen Ocular Genomics, Barcelona, 08028, Spain

**Keywords:** inherited retinal dystrophies, NR2E3, photoreceptor differentiation, retinal development, single-cell RNA-seq

## Abstract

Mutations in *NR2E3* cause two retinal dystrophies with a distinct phenotype. *NR2E3* encodes an orphan nuclear transcription factor that contributes to photoreceptor cell fate determination by repressing cone while activating rod genes. To dissect NR2E3 function, we performed scRNA-seq in the retinas of wild type and two different *Nr2e3* mouse models that show phenotypes similar to patients carrying *NR2E3* mutations. Our results reveal that rod and cone populations are not homogeneous and can be separated into different sub- classes. We identify a previously unreported cone pathway that generates hybrid cones that co-express both cone- and rod-related genes. In mutant retinas, this hybrid cone subpopulation is more abundant, as it includes a subpopulation of rods transitioning towards a cone cell fate. Hybrid photoreceptors with high misexpression of cone- and rod-related genes are prone to regulated necrosis. Overall, our results shed light on the role of NR2E3 in modulating photoreceptor differentiation towards cone and rod fates and explain how mutations in *NR2E3* lead to different visual disorders in humans.

**GRAPHICAL ABSTRACT:** 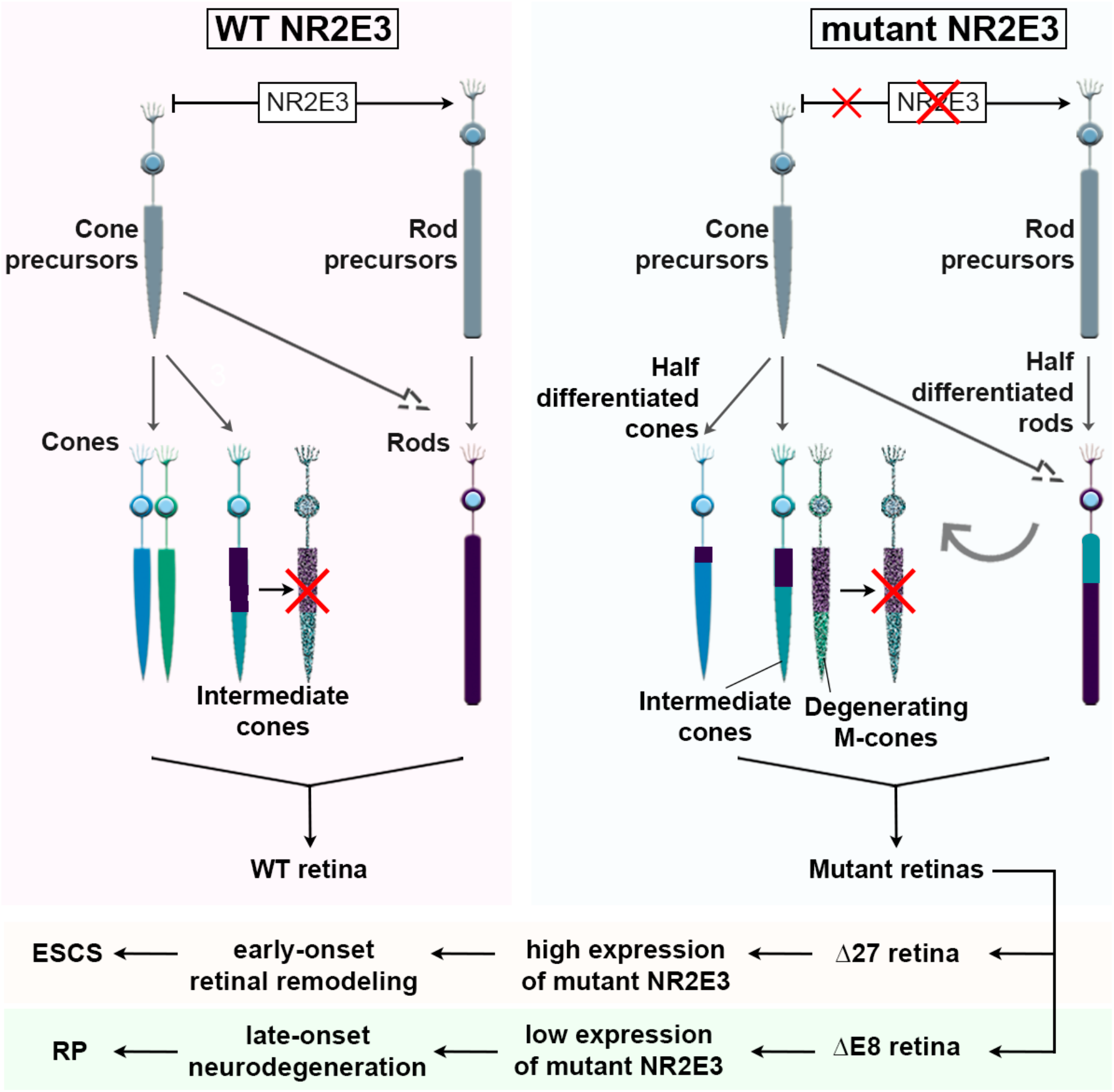

**SYNOPSIS:** Mutations in the gene encoding the retinal transcription factor *NR2E3* cause two different inherited retinal dystrophies: retinitis pigmentosa and enhanced S-cone syndrome.

- scRNA-seq in retinas of wild type and two *Nr2e3* mouse models reveal that rod and cone populations are not homogeneous and can be classified into different sub-classes.
- A previously unreported cone pathway that generates hybrid cones co- expressing both cone- and rod-related genes is identified.
- Hybrid photoreceptors with high misexpression of cone- and rod-related genes are prone to regulated necrosis.
- Expression of rod and cone signature genes change in response to different *NR2E3* mutations thus providing a frame to understand the molecular basis of distinct NR2E3-associated diseases.

## INTRODUCTION

The retina is the light-sensitive tissue responsible for visual perception. Retinal photoreceptors are specialized neurons that detect light photons and initiate the signal transduction from the retina to the brain. Human retina photoreceptors include rods –responsive to dim-light conditions– and three types of cones expressing opsins sensitive to short, medium and long-wavelength light (S-, M- and L-cones, respectively), which mediate colour vision and visual acuity (Nathans et al., 1986; Schnapf et al., 1987). Instead, the mouse retina is formed by rods and cones that express S- and M-opsins. A careful orchestration of regulatory transcription factors is required to determine photoreceptor fate and differentiation during retinal development.

In the developing mouse retina, differentiation into S-cones is the default fate path for photoreceptor precursor cells (PPCs) unless additional regulatory signals determine them towards rod or M-cone identities (Swaroop et al., 2010). In this process, the retinal transcription factor NRL acts upstream of NR2E3, the latter displaying a dual function as both transcriptional activator and repressor in photoreceptors. NR2E3 is an orphan nuclear receptor that can homodimerize and form a transcription regulatory complex together with CRX and NRL to repress the cone-default pathway and activate the expression of rod-specific genes (Chen et al., 2005; Cheng et al., 2006; Peng et al., 2005). In the mature retina, NR2E3 is also necessary for rod homeostasis and maintenance (Cheng et al., 2004; Haider, 2001). Two isoforms of NR2E3 have been identified in both humans and mice (Aísa-Marín et al., 2020a): The long isoform produces the conventional NR2E3 protein (NP_055064.1) whereas the short isoform lacks exon 8 (NP_057430.1), and hence lacks the dimerization and repressor domains. The production of the short NR2E2 isoform appears to be developmentally regulated (Aísa-Marín et al., 2020a), but its physiological function remains to be elucidated.

Diseases associated with *NR2E3* include retinitis pigmentosa (RP, MIM# 611131) and enhanced S-cone syndrome (ESCS; MIM# 268100), whose most severe affectation is also named as Goldmann-Favre syndrome (GFS; MIM# 26800)(Bernal et al., 2008; Coppieters et al., 2007; Gire et al., 2007; Haider et al., 2000; Schorderet and Escher, 2009). *NR2E3* mutations that affect rod maintenance and survival in the mature retina cause RP, a neurodegenerative retinal dystrophy characterized by progressive loss of rod photoreceptors followed later by cones. On the other hand, mutations affecting the inhibition of the cone pathway result in ESCS, characterized by an excess of dysfunctional S-cones in detriment of rods. Except for one *NR2E3* mutation associated with autosomal dominant RP (p.G56R), there is neither a clear phenotype-genotype correlation for recessive retinal dystrophies associated with *NR2E3* mutations – scattered throughout the whole gene– nor the molecular mechanisms causing the distinct clinical phenotypes (Schorderet and Escher, 2009).

We previously generated two *Nr2e3* mouse models using CRISPR-Cas9 editing to delete different domains encoded in the last exon (Aísa-Marín et al., 2020b, 2020a). The *Nr2e3* Δ27 model –carrying a homozygous in-frame deletion that ablates the dimerization domain– shows an ESCS-like phenotype, with profound but non-progressive alterations in retinal function (Aísa-Marín et al., 2020a). The *Nr2e3* ΔE8 model –carrying a complete deletion of exon 8 so that only the short isoform is expressed– shows a RP-like phenotype, characterized by an initial mild rod dysfunction that progressively leads to neurodegeneration, eventually affecting other retinal cell types (Aísa-Marín et al., 2020a). Notably, both mutants present cone-rich invaginations in the central retina, similar to the rosettes observed in the *rd7* mouse retina, a natural model of ESCS caused by a LINE transposon insertion that disrupts the *Nr2e3* gene (Akhmedov et al., 2000; Chen et al., 2006; Haider, 2001).

In addition to histological characterization, retinal transcriptomic analyses through bulk RNA-seq have been widely used to identify the molecular mechanisms underlying pathological processes that cause retinal dystrophies (Bales et al., 2018; Liu et al., 2020; Uren et al., 2014). However, subtle transcriptomic differences between differentiated photoreceptors might go undetected, which is critical while studying the NR2E3 opposing roles in rod and cone transcriptional networks. Here we present single-cell RNA-seq results on our *Nr2e3* models retinas by comparing the distinct cell types and photoreceptor sub-populations that make up the wildtype (wt) and *Nr2e3* mutant retinas.

Our results unveil different and undescribed subclusters within cone and rod cell populations, highlight gene expression changes in rod and cone cells and detect photoreceptor cells with a hybrid rod-cone expression pattern in the *Nr2e3* mutants, all in agreement with NR2E3 dual function as transcriptional activator of rod genes and repressor of cone genes.

## RESULTS

### Single-cell RNA-seq of wildtype and *Nr2e3* mutant retinas reveals different cell populations

First, we sampled whole retinas from wt and mutant mice at stage P40-P80, a timepoint at which Δ27 and ΔE8 mutants differ in their electrophysiological recordings (Aísa-Marín et al., 2020a). In particular, Δ27 mutants are visually impaired due to retinal developmental alterations (ESCS-like phenotype), whereas ΔE8 mutants still show normal electrophysiological responses, as the late-onset retinal degeneration has not yet affected their visual function (RP-like phenotype) (Aísa-Marín et al., 2020a). Retinal samples were subject to droplet-based single-cell RNA-sequencing (scRNA-seq, 10X Genomics Chromium platform), which resulted in the generation of gene expression libraries for a total of 107,628 cells after quality-control filtering. To prevent RNA ambient contamination from extremely abundant and fragile rod cells, scRNA data were further filtered using SoupX (Young and Behjati, 2020). Filtered data is provided in Suppl. Data 1.

Next, we took advantage of a recent publicly available scRNA-seq dataset of wt mouse retinas (Norrie et al., 2019) to integrate our data for robust clustering and cell-type identification (see Methods). Pooled scRNA data from all retinas were normalized, and we identified seven different retinal cell type clusters based on the expression of cell type-specific genes (Fig. S1). These included neurogenic retinal progenitor cells (RPCs), rod and cone photoreceptors, post-synaptic cells, rod bipolar cells, glial cells, and retinal pigment epithelium cells (RPE) (Fig. 1A- D). The post-synaptic cell cluster shared markers with multiple retinal cell types, including bipolar, horizontal, amacrine and retinal ganglion cells. Similarly, the glial cells cluster displayed common markers of astrocytes, Müller glia and microglia. Since we aimed to focus on photoreceptor cell fate, we did not further assign specific cell identities within these two clusters. The composition of cell populations across our pooled samples confirmed that the predominant isolated cell were rod photoreceptors (Fig. 1B), which constitute the primary cell type in the retina of both mice and humans (Jeon et al., 1998).

**Figure 1.**
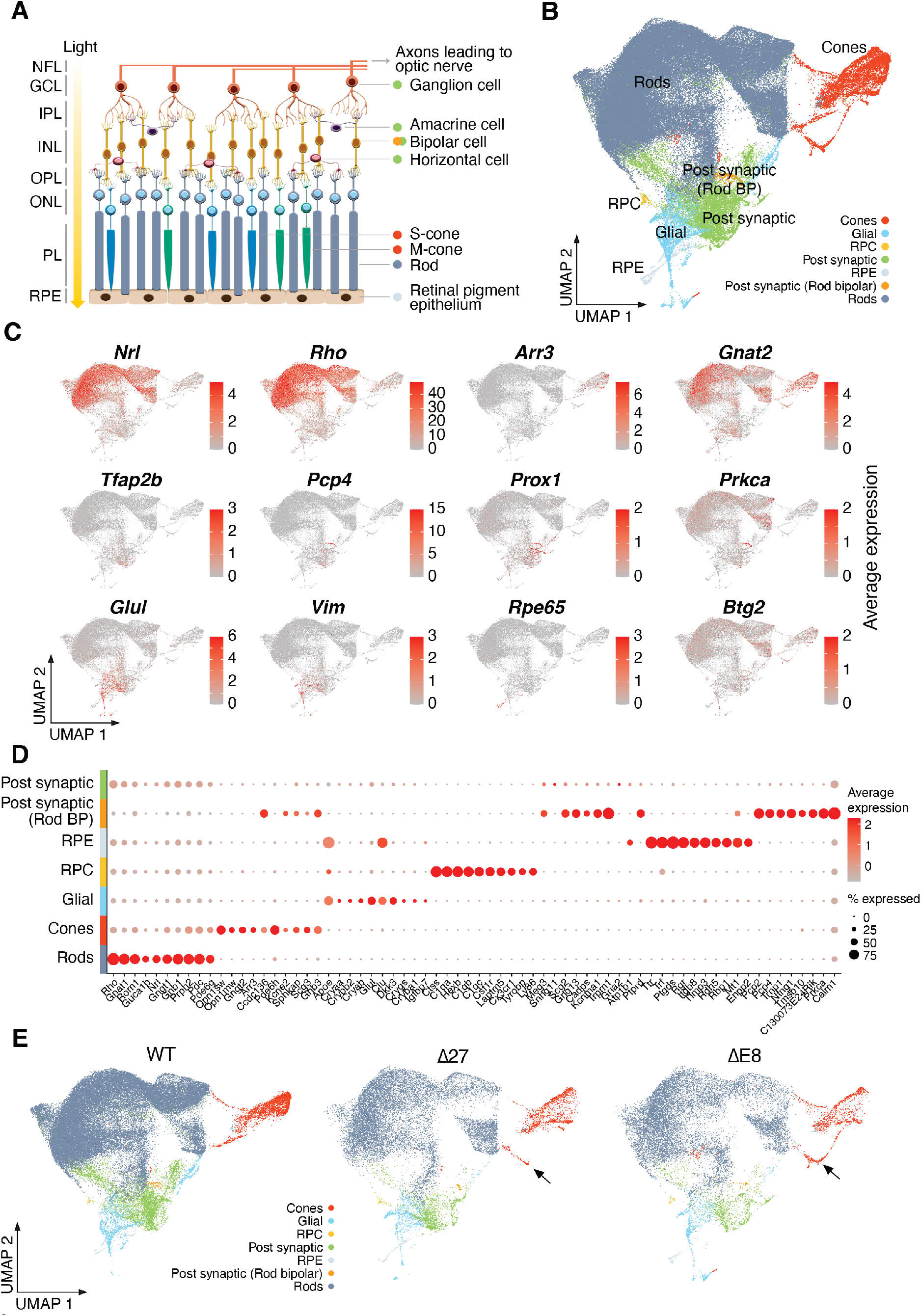
Single-cell RNA-seq of wildtype and Nr2e3 mutant retinas reveals different cell populations. **A.** Scheme shows the different functional cell layers and connections within the mouse retina. Colour dots at the right of each cell type name correspond to the cluster assignment in B. **B.** UMAP plot of sequenced retinal cells showing the clustering of the different cell types. Assignment to main retinal cell types is based on the expression of specific gene markers. **C.** The expression of signature marker genes across cell groups identifies clusters of rod, cone, post-synaptic, rod bipolar, retinal progenitor cells (RPC), glial and retinal pigment epithelium (RPE) cells. **D.** Dot plot of signature genes in the different cell types isolated from mouse retinas. **E.** Differences in the UMAP plot by genotype. Particular subpopulations identified as unique within the cone cluster in the mutant retinas are indicated by a black arrow. See also Figures S1 and S2.

We next calculated the differential expressed (DE) genes for each cluster to identify marker genes whose expression was specific to a particular cell type. High expression levels of *Nrl* and *Rho* identified the rod photoreceptor cluster, and high expression of *Arr3* and *Gnat2* identified the cone cluster (Fig. 1C-1D, Fig. S1). Expression of *Tfap2b* and *Prox1* determined horizontal and amacrine cells (Dyer et al., 2003; Jin et al., 2015; Pérez De Sevilla Müller et al., 2017) among the post-synaptic cell cluster, *Pcp4* determined bipolar(Shekhar et al., 2016) and ganglion cells (Laboissonniere et al., 2019), and *Prkca* specifically defined the rod bipolar cell cluster (Ruether et al., 2010; Woods et al., 2018) (Fig, 1C-D, Fig. S1). The rod bipolar cell cluster was differentiated by the high expression of specific marker genes (Fig. 1D). Expression of *Glul* and *Vim* identified Müller cells (Roesch et al., 2008) and astrocytes (Wunderlich et al., 2015), respectively, in the glial cluster. Finally, high expression levels of *Rpe65* identified a small RPE cell cluster, and *Btg2* was considered a marker of neurogenic RPCs (Trimarchi et al., 2008) (Fig. 1C, Fig. S1). Overall, highly expressed rod genes (*Nrl* and *Rho*) were confined to the rod population of cells (Fig. 1C-D, Fig. S1). These seven major cell type clusters are present in the retinas of all three genotypes (wt, and mutants Δ27 and ΔE8). Notably, the retinas of the two mutants display a new subpopulation of cones that clearly stand aside from the rest of the cells in the cone cluster (Fig. 1E, black arrows).

Next, we examined the expression of *Nr2e3* at the single-cell level. *Nr2e3* was expressed in rod and cone photoreceptors of wt and mutant retinas as expected. Visualization of scRNA-seq reads validated the mutation of each *Nr2e3* mutant model and confirmed the 27-nucleotide in-frame deletion and the complete deletion of exon 8 in Δ27 and ΔE8 mutants respectively (Fig. S2A). The scRNA-seq data confirmed the overexpression of *Nr2e3* in the postnatal Δ27 vs wt retinas, and the low *Nr2e3* expression in the ΔE8 mutant (Fig. S2B), as previously reported using qRT-PCR (Aísa-Marín et al., 2020a). Nonetheless, in all three genotypes, the *Nr2e3* expression ratio between rods and cones was maintained, being 3 times higher in rods than in cones (Fig. S2C). The shallow *Nr2e3* expression in the ΔE8 mutant retinas (which makes this mutant an effectual knockdown) can be partly explained by the reduced stability of the short isoform compared to the long isoform (Fig. S2D).

### *Nr2e3* mutant retinas produce hybrid photoreceptors

To identify more subtle, mutant-specific cell cluster changes in gene expression due to the dysfunction of NR2E3, we calculated the DE genes for the *Nr2e3* mutants compared to the wt samples (Fig. 2A, Suppl. Table 1). Frequent Gene ontology (GO) terms from the DE genes in the rod and cone clusters between the mutants and wt retinas suggest that both cones and rods showed altered gene expression, particularly in genes associated with photoreceptor and neuronal function, but mutant rods show a much higher expression of genes related to neurosensory signalling, phototransduction and photoreception (groups A and B) as well as proteins involved in protein quality control (chaperones, and deubiquitination), autophagy and necrosis pathways (group I) (Fig. 2A, Suppl. Table 1). Consistent with the repressor domain being absent in the mutant NR2E3, most DE genes were overexpressed compared to wt photoreceptors, except with a very clear group of downregulated genes involved in homeostasis, mitochondria and visual longterm conduction and synaptogenesis (Fig. 2A, group G). Opposite alterations in cone versus rod DE can be identified in genes involved in differentiation and eye development, DNA repair mechanisms and translation-associated metabolism (groups C, D and E).

**Figure 2.**
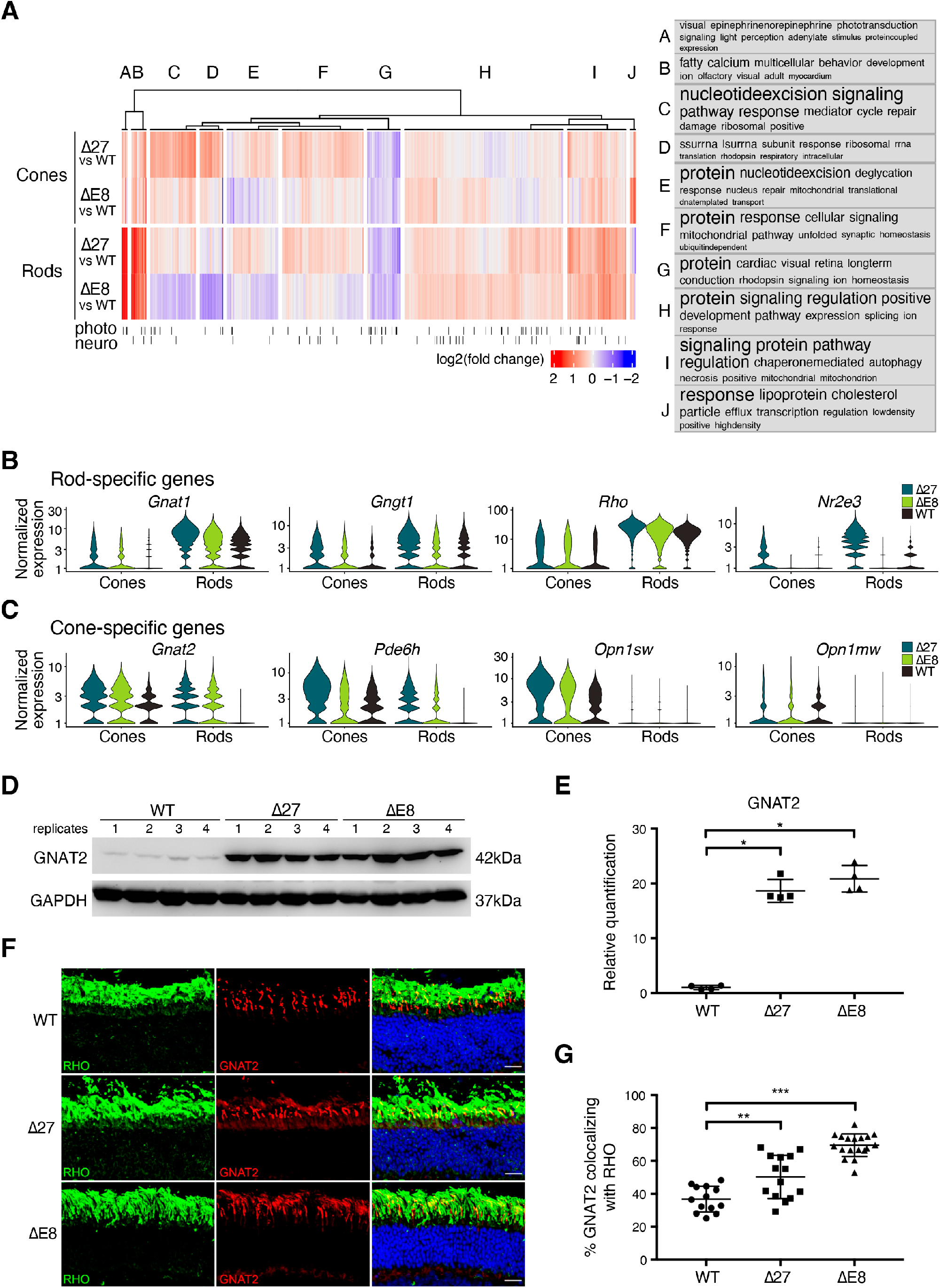
*Nr2e3* mutant retinas produce hybrid photoreceptors. **A**. Log 2-fold change heatmap showing the DE genes sorted by k-means clustering in the rod and cone clusters of Δ27 and ΔE8 mutants vs wt retinas. Frequent biological process GO terms appear in the cloud according to their presence in each cluster. Photo and Neuro indicate specific photoreceptor and neuronal genes, respectively. **B-C**. Expression dysregulation of rod- **(B)** and cone- **(C)** specific genes in the rod and cone populations from mutant Δ27 and ΔE8 vs wt retinas. Note that some rod genes are overexpressed in cones (e.g., *Gnat1, Gngt1*) **(B)**, whereas cone genes are not repressed in rods (e.g. *Gnat2, Pde6h*) **(C)**. See the detailed list in Fig. S3. **D-G.** Confirmation of the overexpression of GNAT2 (cone-specific) in the wt and mutant retinas, by **(D)** immunodetection by western blot of *Nr2e3* wt and mutant retinal lysates shows a 20-fold increase of GNAT2 expression in the mutants. Samples 1-12 are independent biological replicates from the three genotypes (1-4 wt, 5-8 Δ27, 9-12 ΔE8). **E.** Quantification of the data in (**D)**, presented as mean and SD. **F.** Immunohistochemical staining of wt and mutant retinas (n = 3) shows diffuse localization of GNAT2 (in red) in the outer segment of Δ27 and ΔE8 rod photoreceptors (RHO in green). Photoreceptor nuclei are counterstained with DAPI (in blue). **G.** Colocalization of RHO (rod-specific) and GNAT2 (cone-specific) indicative of hybrid photoreceptors is detected in the Δ27 and ΔE8 mutant retinas. Data (from **F**) are represented as mean and SD. See also Fig. S4 and S5.

When focusing specifically on the list of retina-specific genes for light perception and transduction, the altered expression in *Nr2e3* mutant photoreceptors becomes very apparent (Fig. S3). A relevant subset of rod-specific photoreception and phototransduction genes (e.g. *Gnat1, Gntg1, Rho*) are highly and ectopically expressed in the cones of both mutants (Fig. 2B, Fig. S3). Some of these rod-specific genes are also overexpressed in the rods of the mutants (e.g. *Gnat1*, *Gnb1* and *Gngt1*), particularly in the Δ27 samples. In fact, the Δ27 retinas are characterized by a high overexpression of *Nr2e3* in rods and ectopically, in cones (Fig. 2B, more detailed in Fig.S4). Interestingly, mutant cones express *Gngt1*, a rod transducin associated as a marker of foveal cones that is also overexpressed in S-opsin photoreceptors from *Nrl*-null retinal organoids (Kallman et al., 2020; Peng et al., 2019). The overexpression of *Gngt1* and the foveal marker gene *Cyp26a1* in the mutant was confirmed by RT-PCR in total retina samples (Fig. S5).

Moreover, cone-specific genes appear over-expressed because their expression is not correctly repressed. In mutant rods, a subset of genes involved in cone phototransduction (e.g. *Gnat2, Pde6c* and *Pde6h*) are ectopically overexpressed (Fig. 2C, Fig. S3, Fig. S4), but this high ectopic misexpression does not affect all cone genes (e.g. *Opn1mw* and *Arr3*). In addition, in mutant cones most cone-specific genes (such as *Gnat2*, *Pde6h*, *Opn1sw*) are also overexpressed, this increased expression is again more apparent in the Δ27 mutant. Altogether, Δ27 mutants at a young age (the timepoint when the samples were obtained) show a higher number of DE genes than ΔE8 mutants, probably related to the fact that alterations initiate first in the developmentally affected Δ27 model and later in the ΔE8 model, which is affected by a degenerative phenotype (Aísa-Marín et al., 2020a).

In order to experimentally validate these results using a different technique, the overexpression of the cone *Gnat2* marker in mutant retinas was analysed by immunohistochemistry and western blot immunodetection. A significant increase of GNAT2 levels (around 20-fold) was detected in the mutant retinas, in agreement with its upregulation in rod and cone photoreceptors (Fig. 2D-E). In the wt retina, GNAT2 expression is restricted to cones, in contrast to the mutant retinas, where the expression of GNAT2 is diffuse and localized in the outer rod segments (Fig. 2F). Colocalization of GNAT2 and RHO (rod marker) is highly increased in the mutants, confirming the misexpression of this cone-specific gene in rod photoreceptors (Fig. 2G).

Overall, in the *Nr2e3*-mutant photoreceptors, rods ectopically overexpress cone-specific genes, whereas cones ectopically overexpress rod-specific genes, thus unveiling the presence of intermediate hybrid photoreceptor states in the mutant retinas.

### Distinct subpopulations of cones are found in the *Nr2e3* wildtype and mutant retinas

Interestingly, a careful comparison of the UMAP plot by genotype unveiled that the mutant retinas produce a new subpopulation of atypical cone cells (Fig. 1E, black arrows). These cells group aside the rest of the cells in the same cone cluster, pointing towards alterations in the cone differentiation pathway compared to the wt retinas. To further investigate this subpopulation of cone cells detected in the mutant retinas, we performed clustering analysis on the cone cell sub- population (Suppl. Data 2) and identified five subclusters (numbered 0 to 4, Fig. 3A). The total number of cones was lower in the *Nr2e3* mutant compared to wt retinas (Fig. 3B), and the percentage of cells assigned to each subcluster differed between wt and the mutants (Fig. 3C). For instance, subclusters cone_0_, cone_1_ and cone_2_ are present in both the wt and the mutant samples, although their proportion differs: subcluster cone_0_ is more abundant in the wt, while subclusters cone_1_ and cone_2_ are more abundant in the mutant retinas. Subcluster cone_3_ is exclusively detected in the mutant samples, and subcluster cone_4_ is barely detected in the wt, indicating specific alterations in the cone differentiation pathway between the mutant and wt retinas (Fig. 3B-C).

**Figure 3.**
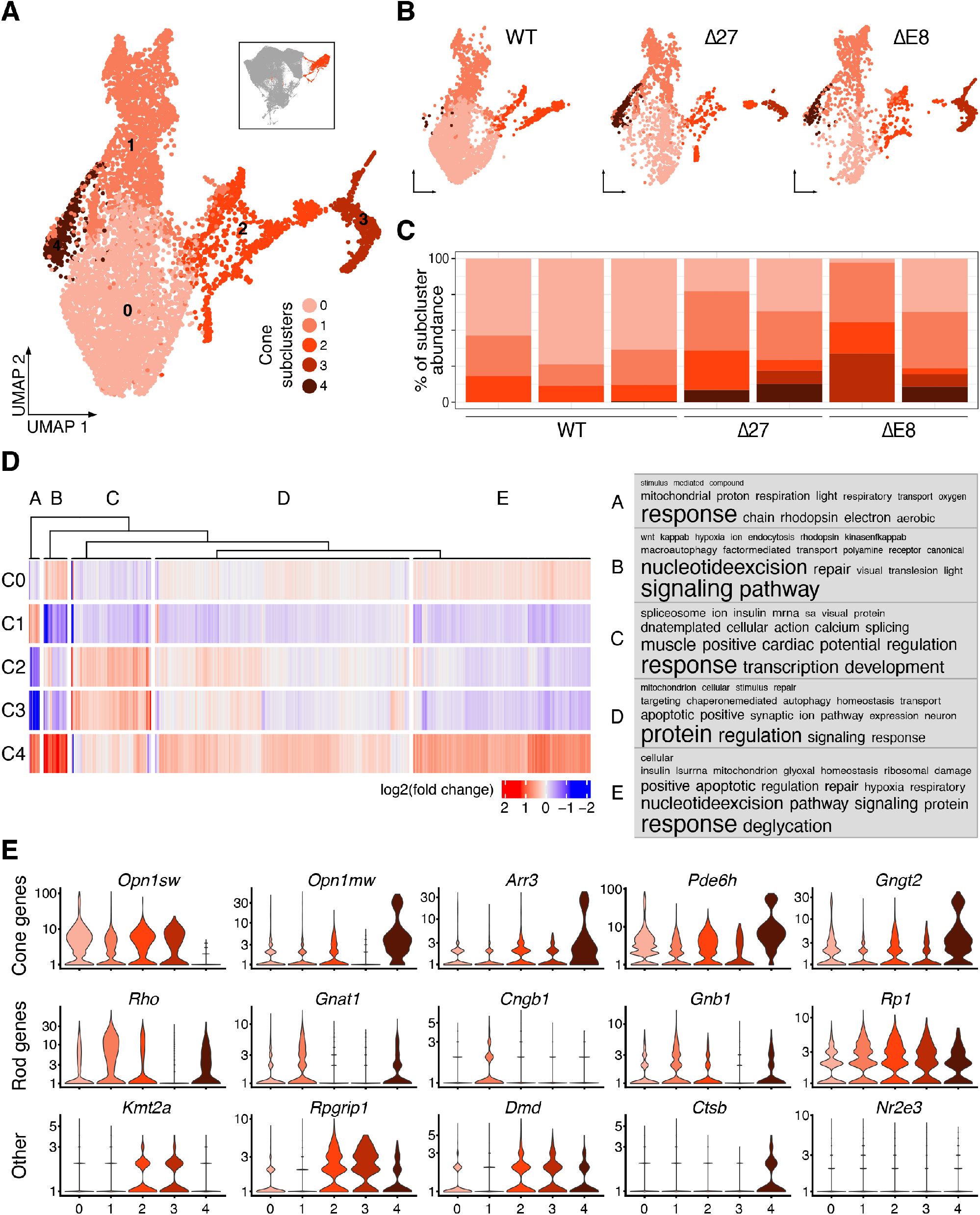
Distinct subpopulations of cones are found in the Nr2e3 wildtype and mutant retinas. **A.** Global UMAP plot of the three genotypes identifying 5 subclusters within the cone population cluster. Each subcluster is indicated with a different shade of salmon-red. **B.** Differences in the UMAP plot by genotype. Note the different burdeos-coloured subcluster at the right in the mutant retinas. **C.** Differences in the percentage of cells in each subcluster per individual. Subcluster cone_0_ is more represented in the wt retina, whereas subcluster cone_1_ is enriched in the mutants. Subclusters cone_3_ and cone_4_ are almost exclusive of the mutant retina samples. **D.** Log 2-fold change heatmap of cone subcluster marker genes sorted by k-means clustering. Frequent biological process GO terms appear in the word cloud according to their cluster. **E.** Identification and assignment to different cone subtypes are based on the expression of specific markers. Subcluster 0 overexpress differentiated cone genes, whereas subcluster cone_1_ overexpresses rod genes. Subclusters cone_2_ and cone_3_ overexpress genes related to retinal development, and subcluster cone_4_, which correspond to M-cones in the *Nr2e3* mutants, also expresses markers associated with neuronal degeneration and cell death. See also Figure S6.

Frequent GO terms of the marker genes in each cone subcluster revealed terms relevant to photoreceptor function (Fig. 3D). Subcluster cone_0_ –with a higher expression of genes necessary for cone phototransduction (*Opn1sw, Arr3, Pde6h, Gngt2*)– represents a subpopulation of fully functional and differentiated cones, while subcluster cone_1_ represents a population of hybrid cones, with both lower expression of cone-specific and concomitant expression of rod-specific genes (*Rho, Gnat1, Cngb1, Gnb1, Rp1*) (Fig. 3E and Fig. S6). The decrease in the number of cells in cluster cone_0_ together with the high increase in cells from subcluster cone_1_ in mutant retinas (Fig. 3C) suggests that hybrid photoreceptors are increased in the mutant retinas in detriment of functional cones compared to wt. This enrichment in cells of subcluster cone_1_ may account for the higher expression of rod genes in the overall cone cell population of the *Nr2e3* mutant retinas.

Cells of subclusters cone_2_ and cone_3_ share the expression of a group of genes involved in retina development that are not expressed in subclusters cone_0_ and cone_1_, including *Kmt2a*, *Rpgrip1* and *Dmd* (Fig. 3E and Fig. S6) (Brightman et al., 2018; Persiconi et al., 2020; Sun et al., 2022). We considered that subcluster cone_3_ (exclusive of the mutants) is derived from subcluster cone_2_: they both correspond to photoreceptor precursor cells (PPCs) already determined to the default cone fate (since they express high levels of S-opsin), but whose transcription signature differs due to NR2E3 misfunction and/or misexpression (Fig. 3E).

Finally, cells from subcluster cone_4_ –mostly restricted to the mutant retinas– are likely degenerating in response to stress, as shown by apoptosis gene expression in GO term analysis (Fig. 3D) and high expression of the necrosis-associated gene *Cathepsin B* (Fig. 3E). One ΔE8 sample does not present subcluster cone_4_ (Fig. 3C), probably indicative of a more severe phenotype, in which all these cones might have already degenerated. Remarkably, all these stressed cones are M-cones (Fig. 3E) that also show high ectopic expression of rod genes (including *Nr2e3*). These traits are also shared with the handful of cone_4_ cells detected in the wt (Fig. S6).

To sum up, single-cell transcriptomic analysis revealed a non-uniform cone population in wt retinas, conformed by a continuum of cells in three main stages, with a high proportion of differentiated cones (subcluster cone_0_, average of 70% of all analysed cone cells), around 20% of halfway differentiated cone cells (subcluster cone_1_), and a low number of precursor cones (subcluster cone_2_, 10%). In contrast, in the *Nr2e3*-mutant retinas the ratios appear to have shifted: the proportion of fully differentiated cones is around 20%, halfway differentiated cones represent 40-50%, and cone cells in an early stage of differentiation (subclusters cone_2_ and cone_3_) account from 15% to 50% of all cones, indicating highly dysfunctional cone subpopulations. Finally, subcluster cone_4_ (M-opsin cones) might include the first cones to degenerate in the *Nr2e3* mutant retinas.

### RNA velocity analysis unveils a population of rods in mutant retinas “transitioning” towards the cone cluster

Similarly to the cone cell population analysis, a subclustering of rod cells (Suppl. Data 3) identified eight subpopulations (Fig. 4A). Despite the significant difference in the total number of rod cells (approximately 60% fewer rods in the mutants than in the wt; Fig. 4B-C), the relative percentage of cells within each rod subcluster remains somewhat similar between all genotypes (Fig. 4C). In contrast to the specific cone subpopulations associated with the mutant genotypes, we could not detect genotype-specific rod subpopulations.

**Figure 4.**
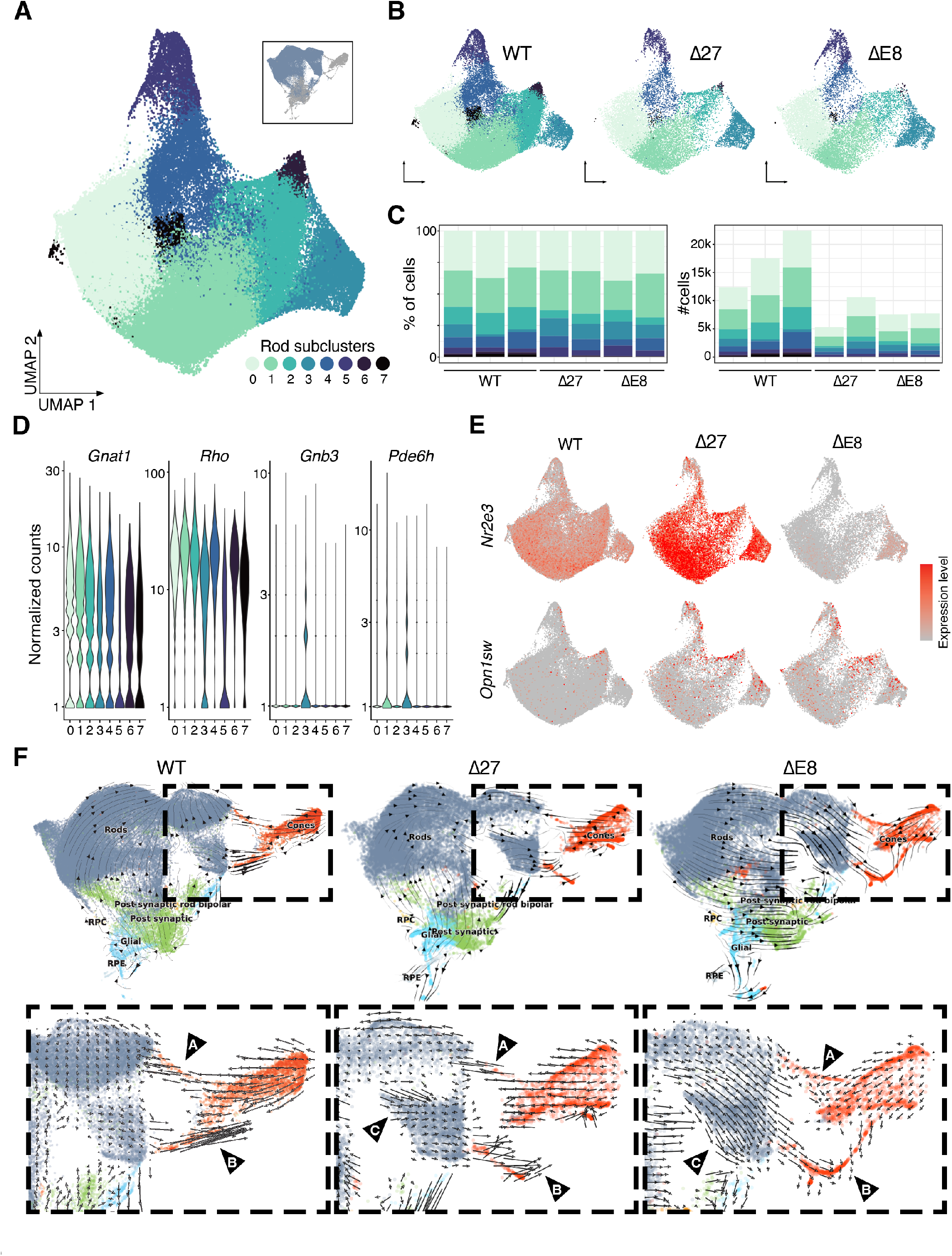
RNA velocity analysis unveils a population of rods in mutant retinas “transitioning” towards the cone cluster. **A.** Global UMAP plot of the three genotypes identifying 8 subclusters within the rod population cluster. Each subcluster is indicated with a different shade of green-blue. **B.** Differences in the UMAP plot by genotype show a decrease in the number of rods in the Δ27 and ΔE8 retinas. **C.** The contribution of each rod subtype (percentage) to the main rod cluster is maintained in all genotypes (left). Δ27 and ΔE8 *Nr2e3* mutant retinas show a 2-fold decrease in the number of rods compared to the wt (right). **D.** Expression of rod and cone genes in the rod subclusters reveals a population (subcluster rod_5_) with low expression of rod genes. **E.** UMAP plot by genotype shows the expression of *Nr2e3* and *S-opsin* in the rod subclusters. *Nr2e3* is upregulated in the Δ27 rods and downregulated in the ΔE8 rods. Of note, in ΔE8, only the subcluster rod_3_ shows the expression of the *Nr2e3* short isoform. Expression of *S-opsin* is only detected in mutant rods, mainly in subcluster rod_5_. **F.** RNA velocity analysis identifies a subpopulation of cones showing strong flow towards the rod cluster in the three genotypes (arrow A); a subpopulation of cones –concurring with the PPCs– transitioning to fully differentiated cones in the three genotypes (arrow B); and a population of rods, which is differentiating towards cones, or tends to acquire cone features (arrows directed towards the cone cluster), exclusively in the mutants (arrow C). See also Figures S7 and S8.

Highly expressed marker genes from subcluster rod_0_ contain frequent GO terms associated with ribosomal and mitochondrially functions, indicative of highly metabolic populations in the retina (Fig. S7A) (Lidgerwood et al., 2021), which require ribosomal activities for protein synthesis and mitochondria for energy. On the other hand, marker genes from other rod subclusters contain GO terms associated with other pathways unrelated to metabolism, indicating two main populations of rods differing in their metabolic activity (Fig. S7A).

Rod-specific genes (*Gnat1, Rho*) were detected in all 8-rod subclusters, although this expression is decreased in subcluster rod_5_ (Fig. 4D, Fig. S7B). The misexpression of cone-specific genes (*Gnb3, Pde6h, Gnat2*) in nearly all subclusters is only detected in mutant retinas, although it is more prominent for the Δ27 samples (Fig. S7B). As expected, *Nr2e3* expression was increased in the Δ27 rods and decreased in the ΔE8 rods, in which we only see expression (corresponding to the *Nr2e3* short isoform) in subcluster rod_3_ (Fig. 4E). We observed an increase in S-opsin expression (*Opn1sw*) in subcluster rod_5_, a cluster characterized by a lower expression of rod genes than the other rod subpopulations (Fig. 4E), which suggests that this subcluster contains rods that are not fully differentiated.

Note that while scRNA-seq provides static information of cellular states at a point in time it does not directly inform about the dynamic transcriptional processes taking part in the different cell populations. We thus used RNA velocity analysis to determine and compare the gene-splicing maturation dynamics in cell populations between the different retinas. In adult retinas, and common to all genotypes, we observe a subpopulation of S-cones transitioning towards the rod cluster (Fig. 4F arrowhead A, Fig. S8), in agreement with the known population of rods that derive from S-cone progenitors during mouse retina development (Kim et al., 2016). Besides, the subpopulation of cones previously identified as PPCs (subclusters cone_2_ and cone_3_) show a strong directional flow towards the differentiated cones, confirming our previous assignment (Fig. 3, Fig. 4F, arrowhead B).

Comparing the results of the RNA velocity analysis of wt and mutant retinas, we identified a population of rods in both mutants showing a strong directional flow towards the cone cluster. This trend is higher for the ΔE8 (Fig. 4F, arrowhead C), indicating that this rod subpopulation could be “transitioning” towards the canonical cone pathway. Another relevant difference, particularly in the ΔE8 mutant, is the detection of strong RNA dynamic flows within each main cell type population towards the cone cluster.

### Retinal remodelling processes in post-synaptic and glial cells occur in response to photoreceptor degeneration

Due to the role of NR2E3 in photoreceptor cell fate determination, NR2E3 misfunction was expected to cause a larger impact on the photoreceptor cell population. Nonetheless, we also analysed potential alterations in other main retinal cell types (Fig. 5A). The RPE cluster only presented a small number of DE genes (Fig. 5B), whereas glial, rod bipolar and the progenitor cell (RPC) clusters showed more relevant changes in gene expression in both mutants, and these alterations affected genes involved in different pathways (Fig. 5C). For instance, rod bipolar cells showed underexpression of genes involved in oxidative stress and DNA repair pathways in contrast to the higher expression of this set of genes in glial cells (Fig. 5A, group B). These changes are likely to be secondary to photoreceptor alteration since *Nr2e3* is not reported to be expressed in these cell types. As occurred in the rod and cone cell populations, retinal cells from the Δ27 mutant showed a higher number of DE genes compared to those of ΔE8 (Fig. 5B). For instance, Δ27 retinas show a 1.5-fold increase in the number of DE genes in the glia and rod bipolar clusters compared to ΔE8, whereas in rod bipolar cells, for some genes, the Δ27 mutants show a 15-fold increase (more than one order of magnitude) compared to ΔE8 (Fig. 5B). The high transcriptomic alteration of the Δ27-rod bipolar cells may explain the decreased functionality observed in the whole retina of Δ27 mutants, not yet detected in the young adult ΔE8 mutants even though they have similar –although less pronounced– changes in photoreceptor cells.

**Figure 5.**
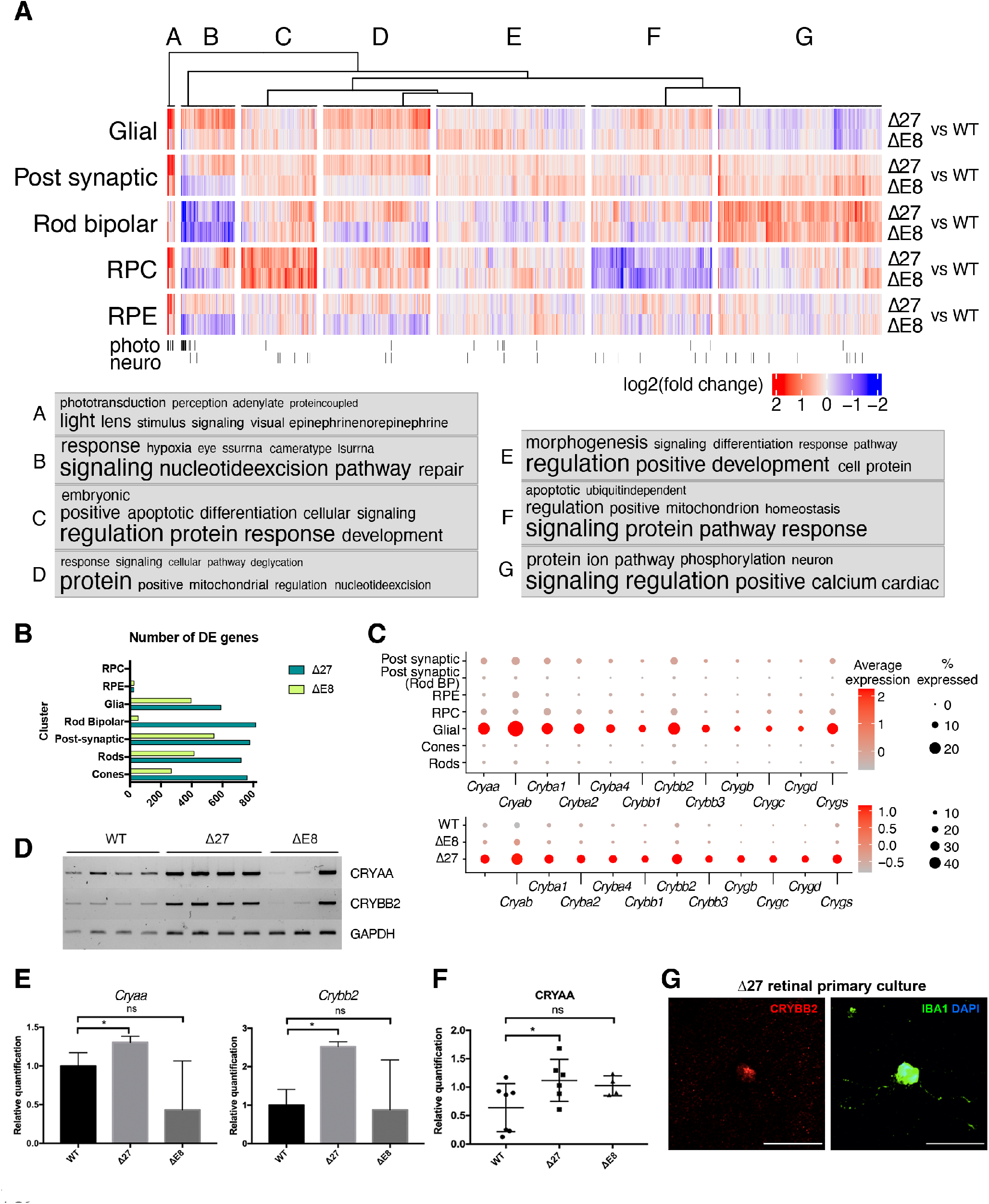
Altered gene expression in *Nr2e3* mutant photoreceptors impact in other neuronal layers and indicate extensive retinal remodelling processes affecting glial, bipolar and progenitor cells. **A.** Log 2-fold change heatmap of DE genes in different retinal cell types in Δ27 and ΔE8 mutants vs wt retinas sorted by k-means clustering. Frequent biological process GO terms appear in the cloud according to their presence in each cluster. Photo and Neuro indicate specific photoreceptor and neuronal genes, respectively. **B.** Number of DE genes in the mutant vs wt retinas in the seven main cell type clusters. Note that the number of DE genes is higher in the Δ27 retinas compared to the ΔE8 retinas. **C.** Dot plot showing average and percentage expression of several genes of the crystallin (cry) family in the seven main cell type clusters (upper panel). The expression of Cry genes, indicative of retinal stress, is increased in the glial cell cluster in the Δ27 mutants (bottom panel). D. *Cryaa* and *Crybb2* overexpression were validated by RT-PCR and Western Blot **(E, F)** of retinal lysates. Samples 1 to 11 D correspond to independent biological replicates from the three genotypes (1-4 wt, 5-8 Δ27, 9-11 ΔE8). Data are represented as mean and SD**. G.** Immunocytochemistry of Δ27 primary retinal cultures show co-expression of CRYBB2 (red) and IBA1 (staining microglia, green) in the same cell, confirming the expression of cry genes indicative of stress-associated remodelling in the glia.

High over-expression of crystallin (Cry) genes was also detected in the glial cell cluster of the Δ27 compared to wt or ΔE8 retinas (Fig. 5A group A, Fig. 5C). Comprising two families, α- and βγ-crystallins are the most prevalent proteins in the lens. Although their precise function out of the lens has yet to be determined, crystallins have been proposed to act as stress proteins in the retina and are considered markers of retinal remodelling (Andley, 2007; Templeton et al., 2013). Overexpression of *Cryaa* and *Crybb2*, belonging to the two crystallin families, was confirmed by RT-PCR (Fig. 5D-E) and Western Blot (Fig. 5F). Colocalization of CRYBB2 and the microglia cell marker IBA1 in Δ27 primary retinal cultures was also observed (Fig. 5G), supporting the hypothesis that the expression of crystallins in glial cells might reflect an induced response to retinal stress.

DE genes between mutants and wt revealed that RPCs in the mutants have lower expression of energy metabolism and mitochondrial genes associated with neuronal function, and overexpression of genes related to differentiation and developmental genes (Fig. 5A, groups F and C, respectively). Additionally, Δ27 retinas frequently show processes associated with anomalous metabolism like metabolic process, glycolysis, regulation of translation and rRNA processing (Fig. 5A groups A, B and D). Overall, these results support that the alterations observed in Δ27 mutant photoreceptors also affect other retinal cell types, particularly rod bipolar and glial cells, which reprogram into abnormal metabolic phenotypes in response to the retinal degeneration process, the so-called retinal remodelling (Jones et al., 2016; Pfeiffer et al., 2020).

### Hybrid photoreceptors degenerate via regulated necrosis

Apoptosis is considered the most common programmed cell pathway. Nonetheless, retinal degeneration does not always occur by apoptosis, and photoreceptor degeneration in Δ27 and ΔE8 mutant retinas was shown not to be caused by this cell death pathway (Aísa-Marín et al., 2020a). Notably, *Ctsb* is particularly upregulated in the subcluster cone_4_, corresponding to degenerating M-cones (Fig. 3E). *Cstb* encodes *Cathepsin B* (CTSB), recently associated with regulated necrosis, a form of non-apoptotic cell death (Berghe et al., 2014; Kuang et al., 2020). To assess whether neurodegeneration of M-cones in the *Nr2e3* mutants might be due to CTSB-related necrosis, the expression and localization of CTSB were analysed by immunohistochemistry in retinal sections (Fig. 6A). Our results confirmed that CTSB was preferentially expressed in M-cones compared to S-cones and that CTSB expression was increased in the mutant retinas (Fig. 6B). Interestingly, CTSB was highly expressed in the cone-rich invaginations characteristic of the *Nr2e3* mutant retinas, suggesting that the early onset degeneration is initiated in these specific areas due to necrosis (Fig. 6A).

**Figure 6.**
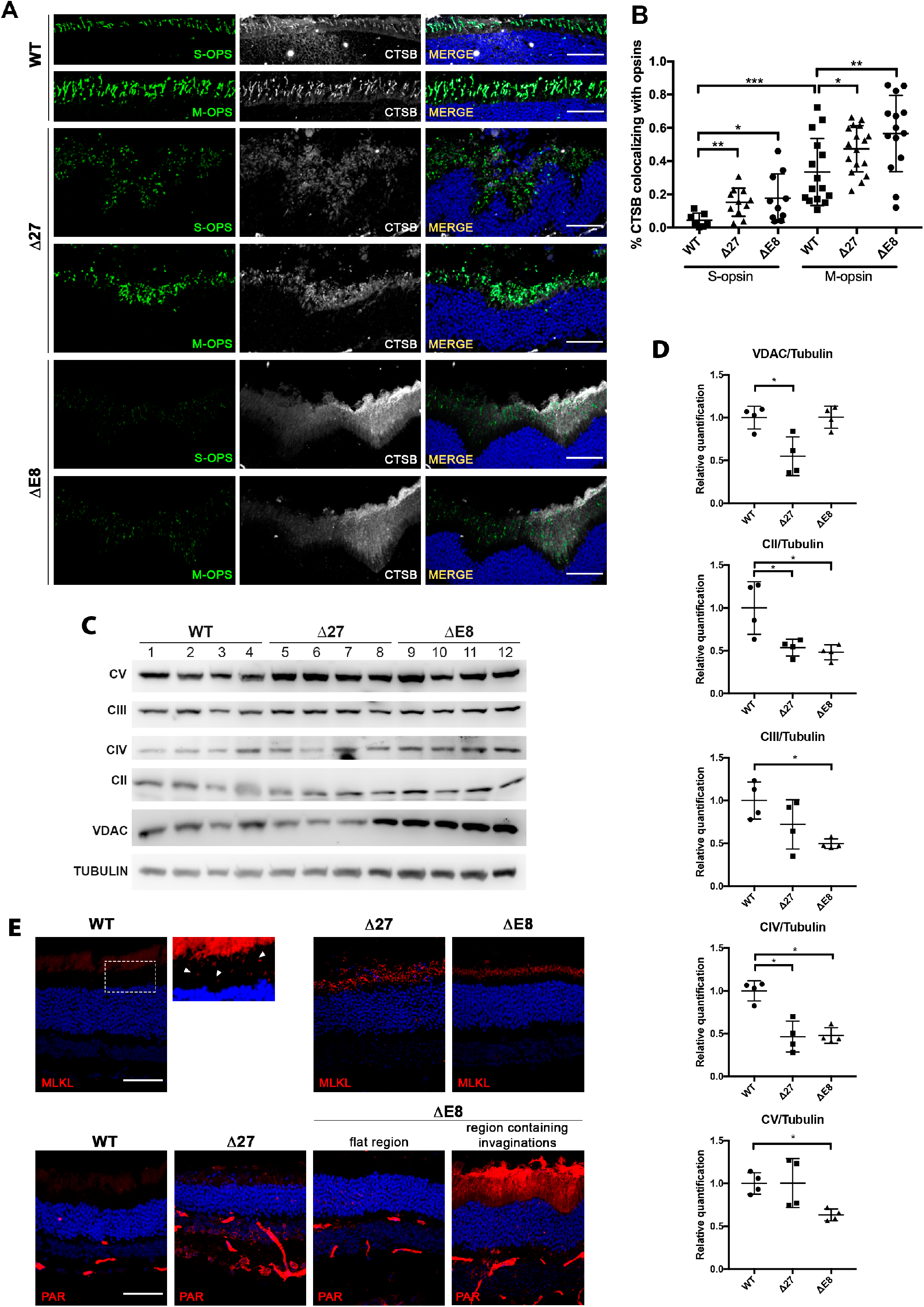
Hybrid photoreceptors degenerate via regulated necrosis. **A.** Cathepsin B (CSTB, associated with regulated necrosis) immunostaining together with (**B**) colocalization with M- and S-opsins (in green) reveals increased CTSB expression in the M-cones compared to the S-cones in all genotypes, and increased CSTB expression in the mutant retinas compared to wt. Note the higher CSTB expression in the cone-rich invaginations in the mutant retinas. Samples 1 to 12 correspond to independent biological replicates from the three genotypes (1-4 wt, 5-8 Δ27, 9-12 ΔE8). Data are represented as mean and SD. **C.** Immunodetection and **(D)** quantification of the components of the mitochondrial OXPHOS complex reveals a reduced mitochondrial mass (VDAC/tubulin) in the Δ27 retinas and lower levels of mitochondrial complex proteins, particularly Complex II and Complex IV (CII and CIV), in the retinas of the two models, indicating mitochondrial alterations in the mutant retinas. Data are represented as mean and SD. **E.** ph-MLKL and PAR immunostaining, respectively indicative of the two necrotic pathways necroptosis and parthanatos, reveal an increased signal in the Δ27 and ΔE8 mutant retinas compared to wt, pointing to retinal cell death in the mutant models by alternative pathways other than apoptosis.

Regulated necrosis is activated by impaired oxidative phosphorylation (OXPHOS) in the mitochondria (Fatokun et al., 2014; Koo et al., 2015; Murata et al., 2019; Nagley et al., 2010). As transcriptional alteration of mitochondrial genes (altered pathways include energy metabolism and mitochondrion homeostasis) were detected in the *Nr2e3* mutant photoreceptors (Fig. 2A, Suppl. Table 1), we sought to confirm the misexpression of mitochondrial proteins in our mutants. The Δ27 mutant retinas showed a statistically significant decrease in VDAC expression (Fig. 6C-D), indicating mitochondrial mass reduction. Downregulation of multiple components of the OXPHOS chain, associated with impaired mitochondrial metabolism, was also observed in the Δ27 and ΔE8 retinas (Fig. 6C-D). These results confirm that the mutant retinas possess an altered mitochondrial metabolism, with reduced mitochondrial mass in the Δ27 retinas and a lower number of functional mitochondrial complexes in the ΔE8 retinas. This suggests an impaired mitochondrial respiration as a plausible trigger for regulated necrosis.

These findings prompted us to analyse other specific necrotic mechanisms –such as necroptosis and parthanatos– that could be additionally involved in photoreceptor degeneration in the mutant retinas (Fig. 6E). Immunodetection of the necroptotic pathway induced via phosphorylated MLKL (ph-MLKL) in *Nr2e3* wt and mutant retinas showed differences between the genotypes. In the wt retina, ph-MLKL was detected in small puncta in the cytoplasm, where it is reported to associate with endosomes and assist in endosomal transport and vesicle release (Yoon et al., 2017). In the Δ27 and ΔE8 mutant retinas, MLKL was aggregated into larger cytoplasmic clusters that accumulated at the plasma membrane (Fig. 6E), where they have been described to compromise membrane integrity and cause cell death (Samson et al., 2020), thus confirming the activation of the necroptotic pathway.

On the other hand, parthanatos is a form of programmed cell death characterized by the overactivation of PARP-1, which uses NAD+ and ATP to synthesize PAR (Fatokun et al., 2014). Therefore, PAR accumulation is commonly used to detect parthanatos. In the Δ27 mutant retinas, there was an increase in the PAR staining in the inner segment of photoreceptors, inner nuclear (INL) and ganglion cell layers (GCL) (Fig. 6E). In contrast, the ΔE8 mutant retinas showed PAR levels comparable to those of the wt in the flat (without invaginations) regions of the retina compared to a highly increased signal in the cone-rich invaginations, reinforcing the findings that degeneration initiates in these regions.

Therefore, retinal neurodegeneration in the retinas of our *Nr2e3* mutants does not occur by apoptosis but rather several alternative necrotic pathways are activated, such as necroptosis and parthanatos, leading to photoreceptor death.

## DISCUSSION

*Nr2e3* mutants display the same seven major cell types identified in the wt retinas. However, both types of photoreceptors –rods and cones– from the mutant retinas show misregulation of rod and cone phototransduction genes. As the Δ27 and ΔE8 mutants lack a functional long isoform of *Nr2e3* –which encodes the dimerization and repressor domains– we surmised that the role of NR2E3 as a transcriptional repressor of cone-specific gene expression was impaired in the mutants and that only the transcriptional activator function remained. This is indeed the case since the expression of the cone pathway genes cannot be fully repressed and are overexpressed in the rod and cone clusters. Rod genes are also overexpressed in the rod and cone clusters since all the NR2E3 protein expressed by photoreceptors could only act as a transactivating transcription factor. The upregulation of rod genes is more notable in the Δ27 mutant, agreeing with the high overexpression of *Nr2e3* in the Δ27 compared to the ΔE8 mutant, which is more akin to a knockdown *Nr2e3* model. Misregulation of phototransduction genes generates hybrid or intermediate photoreceptors, in which both rod- and cone-specific genes are expressed in the same cell type.

Although NRL is the main transcription factor for rod differentiation, NR2E3 is required to secure and maintain rod photoreceptor commitment and homeostasis. Therefore, a substantial decrease in the number of rods is detected in the *Nr2e3* mutant compared to the wt retina. Several reasons can explain this decrease in the rod number: (i) a global reduction in the number of rods generated during the development (at the expense of an increased number of cones), (ii) the misfunction and subsequent degeneration of the existent rods, or (iii) a combination of both.

During mammalian retina development, photoreceptor precursors (PPCs) follow a default pathway to differentiate into S-cones unless other regulatory signals direct them to the rod or M-cone identity (Swaroop et al., 2010). In addition to rods originating from rod precursors, some mature rods derive from a pool of PPCs initially expressing S-opsin, which redirect their fate towards the rod pathway, demonstrating the plasticity of initial S-cone photoreceptors (Kim et al., 2016) (graphic model in Fig. 7A). Some authors hypothesize that the increase in the number of S-cones observed in the *Nr2e3* mouse mutants comes from the fraction of rods derived from the default S-cones that fail to fully differentiate into rods due to lack of functional NR2E3 (Coppieters et al., 2007; Xie et al., 2019). The RNA velocity analysis in our retina samples detects a strong flow from the S-cone population to the rod cluster in both the wt and the mutants, consistent with the subpopulation of default S-cones differentiating into rods. Nonetheless, we also observe a transition flow from the rod population towards the cone cluster in the mutants that is not present in the wt (Fig. 4F). Altogether, our results indicate that the increased number of S-cones observed in the Δ27 and ΔE8 mutants arise from photoreceptors committed to the rod pathway that transition towards cone fate instead of arising from the particular population of rods derived from PPCs expressing S-opsin (model in Fig. 7B).

**Figure 7.**
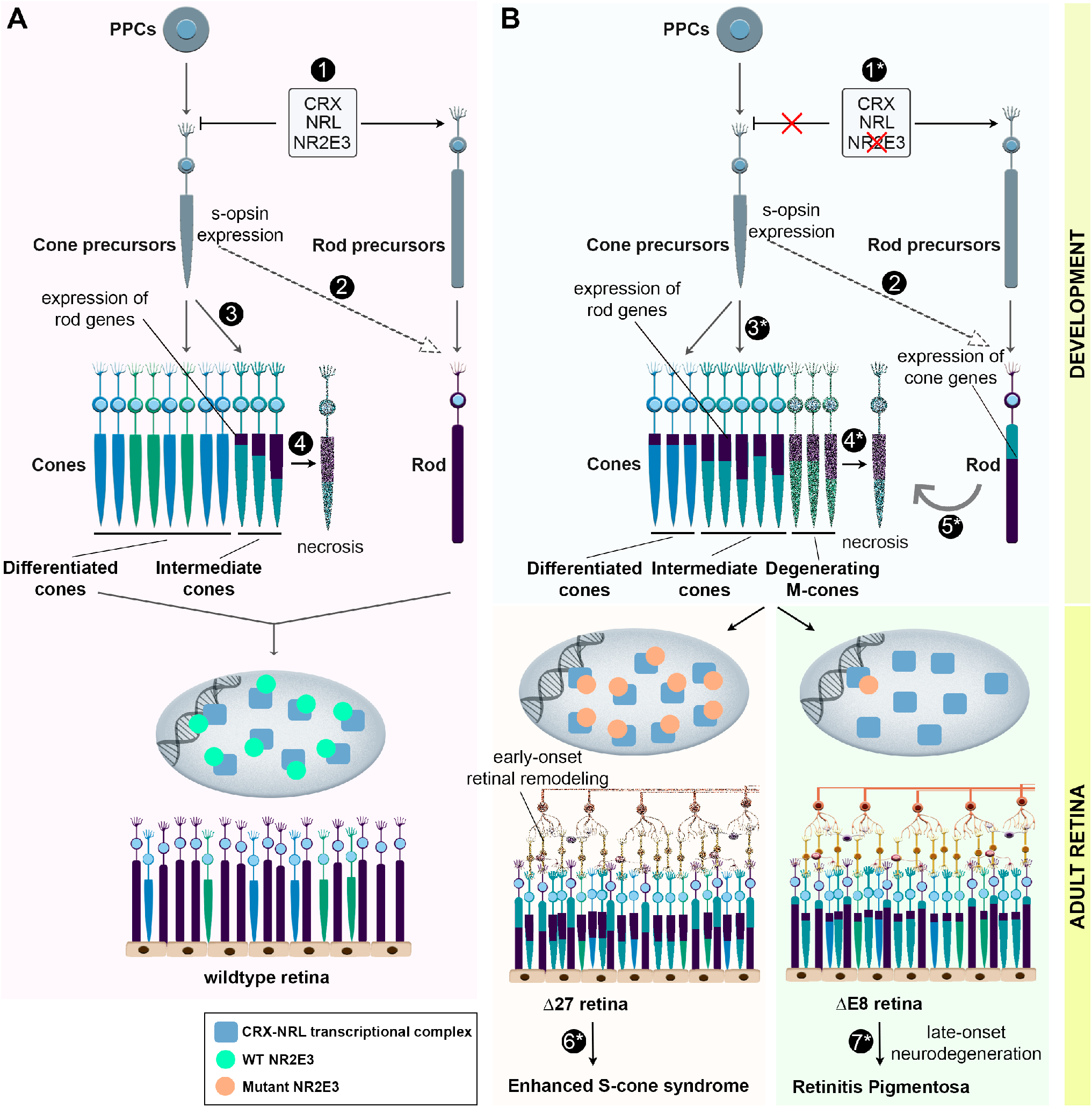
Proposed model for NR2E3-associated disease mechanisms. **A.** During the development of the wt retina, NR2E3 contributes –together with CRX and NRL– to both repression of the cone default pathway and activation of rod genes, thus committing photoreceptors towards the rod fate (1). A fraction of cone precursors expressing S-opsin also contribute to the rod population (2), while most cone precursors differentiate into S- or M-cones. The commitment to rod or cone fate is not fully mutually exclusive, and a small number of cones highly express both cone and rod genes (3). These intermediate cones degenerate via regulated necrosis (4). **B.** Loss or misfunction of NR2E3 due to mutations impairs securing the commitment to the rod fate; repression of cone genes partially fails while rod genes are activated, thus resulting in a population of hybrid rods (1*). The fraction of cone precursors that contribute to the rod population is maintained (2). However, the largest pool of the cone population fails to fully differentiate and become intermediate cones displaying high expression of both cone and rod genes (3*). Mutant intermediate M-cones that highly express rod genes degenerate and die via regulated necrosis (4*). Besides, in the mutant retinas, a fraction of hybrid rods transition backwards to a more cone-like state (5*). In the Δ27 retinas, with a similar phenotype to human ECSC, overexpression of the mutant NR2E3 may cause a dominant transcription factor negative effect that results in higher misexpression of rod and cone genes, which leads to retina remodelling. Some residual NR2E3 function remains and retinal degeneration proceeds more slowly (6*). In contrast, in the ΔE8 retinas, the expression of mutant NR2E3 is very low and the effect on misexpression of rod and cone genes is not as pronounced. However, the lack of NR2E3 affects photoreceptor homeostasis and results in the progressive neurodegenerative phenotype characteristic of RP (7*).

One of the main conclusions from our single-cell retinal analysis is that cones and rods are not homogeneous populations of photoreceptor cells but instead present a continuum of differential phenotypes. The use of marker genes allowed the identification of these different subpopulations within the large heterogeneous rod and cone clusters. Other authors had already identified rod subpopulations that differ in their direct synaptic contacts (Tsukamoto et al., 2001). Our data further dissects the large rod population by identifying a subpopulation of rods characterized by high metabolic requirements, as indicated by increased expression of ribosomal and mitochondrial genes. The presence of distinct rod subpopulations might result from specific adaptations for the different requirements of the rod subtypes.

Interestingly, although the number of rods is significantly decreased in the mutants, the number of subclusters is similar between the three genotypes, and only the percentage of cells in each subcluster varies. These results suggest that mutations in *Nr2e3* cause a shift of cells among rod subpopulations, rather than adding or subtracting specific subpopulations. The similar number of subclusters can be explained if: (i) all subtypes of rods degenerate uniformly, or (ii) rod population may be dynamic, with a flexible modulation between the distinct types of rods. Thus, the rod differentiation pathway is not directly affected by mutations in *Nr2e3*, which reinforces the role of NR2E3 in maintaining the fully differentiated rod fate and favouring rod survival and homeostasis.

Remarkably, *Nr2e3* mutations –at least in young adult retinas– have a larger and more relevant impact on the composition and identity of cone subclusters since the proportions between the different cone subpopulations are inverted, and new subclusters appear as exclusive or nearly exclusive of mutant retinas. The number of fully differentiated cones diminishes whereas subpopulations of intermediate cones or precursor cone cells are much enriched in the mutant retinas, accounting for up to 70% of all cone cells.

Of note, our results strongly support that *Nr2e3* mutations cause selective M-cone degeneration through the necrotic pathway, a common form of non-apoptotic cell death. In fact, there is increasing evidence that alternative cell death mechanisms different from apoptosis play a major role in retinal neurodegeneration (Arango-Gonzalez et al., 2014; Yan et al., 2021). Degeneration seems to start in the regions of the mutant retina presenting the cone-rich invaginations; however, the specific mechanism by which degeneration occurs selectively in M- versus S-cones remains unknown. We hypothesize that the enhanced plasticity observed in S-cones during the development increases its resilience against transcriptional changes compromising the cone identity (e.g., by expression of rod-specific genes), thus causing early-onset degeneration only in the M-cones. ESCS patients show increased S-cone and decreased M- and L-cone sensitivities, although this varies among individuals (Ripamonti et al., 2014). Together with data obtained in zebrafish (Xie et al., 2019), our results suggest that the decreased M- and L-cone sensitivities in ESCS patients might be due to a combination of both, the increased number of S-cones and the selective loss of M- and L-cones, as a consequence of *NR2E3* mutations.

Interestingly, a small number of cones in the wt retinas also show transcription of rod genes. By integrating the scRNA-seq data from the three genotypes (wt, Δ27 and ΔE8) to generate the main clusters and subclusters shown in this study, we were able to detect this tiny population of cones in the wt retinas –which is highly increased in our mutants– that express rod genes and subsequently degenerate (subcluster cone_4_). This fact suggests that the differentiation process towards cone or rod fate is neither strict nor fully mutually exclusive and may fail in some cells.

In response to photoreceptor deterioration, the retina suffers a process of negative plasticity called retinal remodelling, which comprises multiple mechanisms like alterations in retinal metabolism and neuronal network topologies (Jones and Marc, 2005; Jones et al., 2016; Pfeiffer et al., 2020). We hypothesize that retinal remodelling occurs in the *Nr2e3* mutants, as suggested by the frequent GO terms in specific pathways in the rod bipolar and glial cells, particularly in the Δ27 retinas, which also show overexpression of the cry family of genes triggered in response to stress. In fact, all the observed expression alterations are more pronounced in the Δ27 than in the ΔE8 mutants, not only by the increased gene misexpression in rods and cones but also by the higher number of DE genes in all cell types, particularly bipolar cells, highly indicative of extensive retinal remodelling in the Δ27 retinas. Concerning human disease, the Δ27 is a model of the ESCS whereas the ΔE8 retina shows a late-onset RP phenotype (Aísa-Marín et al., 2020a). Our data suggest that the ECSC-like retinal dystrophy detected in the Δ27 retinas at early stages is mostly due to this early-onset retinal remodelling rather than to rod/cone gene misexpression, as young ΔE8 retinas which also show gene misexpression only display slightly reduced responses in rod cells (Aísa-Marín et al., 2020a). Therefore, and according to our proposed model (Fig. 7), a potential dominant negative effect of mutated NR2E3 (as may happen in the Δ27 model) would mostly cause early-onset retinal remodelling, whereas loss of function/knockdown mutations (as in ΔE8) would affect photoreceptor homeostasis and survival at later stages, leading to RP-like progressive photoreceptor attrition.

Overall, our results support that *Nr2e3* misfunction: a) causes transcriptional misregulation of cone and rod marker genes; b) changes photoreceptor fate – particularly evident in cone cell populations– with an enrichment in cones that are not fully differentiated; c) demonstrates that cone and rod populations are not homogeneous but composed of different subpopulations with specific requirements and transcriptional signatures; d) indicates that photoreceptor differentiation is not a strict and mutually exclusive process that generates either rods or cones –even in the wt retinas–, but there is instead a hybrid intermediate photoreceptor subpopulation in a stage that can be directed towards the rod or the cone fate; and e) demonstrates that high overexpression of both rod and cone marker genes in the same cell compromises photoreceptor identity, which will then degenerate through necrotic pathways alternative to apoptosis, such as necroptosis or parthanatos. Depending on the *NR2E3* mutation, photoreceptor fate commitment and/or photoreceptor homeostasis might be differentially affected, thus leading to either the ECSC or the RP phenotype.

Modulation of the NRL/NR2E3 pathway has been recently proven to be effective in restoring retinal integrity and function across various IRD models (Li et al., 2020; Nakamura et al., 2017; Yu et al., 2017). In this context, our study provides the basis to understand how rod and cone populations change in response to NR2E3 function/dysfunction and the potential side effects –such as retinal remodelling– that it might cause in the long term.

## FUNDING

I.A.-M. is recipient of the APIF grant (Universitat de Barcelona), and the Company of Biologists grant for short term mobility. This research was supported by grants PID2019-108578RB-I00 (Ministerio de Ciencia e Inno-vación/FEDER), 2017 SGR738 (Generalitat de Catalunya) to G.M. Work in the Vaquerizas laboratory is supported by the Medical Research Council, UK (award reference MC_UP_1605/10 to J.M.V.), the Academy of Medical Sciences and the Department of Business, Energy and Industrial Strategy (award reference APR3\1017 to J.M.V.).

## ACKNOWLEDGEMENTS

We are grateful to the associations of patients affected by retinal dystrophies for their constant support, particularly Retina Asturias, and Asociación “Muévete por los que no pueden”. We also acknowledge past and present members of our research group for helpful discussions.

## AUTHOR CONTRIBUTIONS

I.A.-M. and Q.R. have performed the experiments, analysed the data; designed figures and written the draft manuscript; N.D. and L.C. have contributed to some figures; J.M.V. and G.M. provided the initial concept and funding, have supervised the work and analysed the data. All authors have revised the manuscript.

## DECLARATION OF INTEREST

The authors declare no conflict of interest.

## STAR METHODS

### Resource Availability

#### Lead contact

Further information and requests for resources and reagents should be directed to Gemma Marfany.

#### Data and code availability

Raw single-cell RNA-seq data have been deposited at ArrayExpress under the accession E-MTAB-12183 and are publicly available as of the date of publication. Accession numbers are also listed in the key resources table. Further information and requests for scRNA-Seq data and analyses should be directed to Juanma Vaquerizas.

### Animals and ethical statement

Animal handling, euthanasia and surgical dissections were performed according to the ARVO statement for the use of animals in ophthalmic and vision research, following the guidelines for animal care of the University of Barcelona and with the approval of the Ethics Committee for animal experimentation File number FUE-2019-00965313, ID 2MDLDY4WZ), and the Bioethics Committee.

### Single-cell dissociation, methanol fixation, rehydration and cDNA library preparation

Two retinas per animal (three wild-type and two per each mutant) were dissected in Neurobasal media and placed on ice. Retinas were transferred to a papain solution (Neural Tissue Dissociation Kit – Postnatal Neurons, Milteny Biotec) and incubated at 37°C for 15 min (agitation at 5-min intervals), following the manufacturer’s instructions. DNase I (10U/ml, Roche) was added and incubated at 37°C for additional 5 min. Samples were homogenized by gently pipetting, and cells were centrifuged at 400 g for 5 min at 4°C. Methanol fixation and rehydration were performed according to the Single Cell RNA Sequencing guidelines of 10X Genomics. Briefly, a minimum of 1x10^6^ cells were resuspended in 200 ml of cold PBS and 800 ml of cold methanol. Cell suspension was placed on ice for 30 min prior to transferring to -80°C for long-term storage for up to 6 weeks. For rehydration, cells were placed on ice for 5 min to equilibrate them to 4°C. Cells were centrifuged at 1000 g for 5 min at 4°C and resuspended in 300 ml of Wash Resuspension Buffer (0.04% BSA, 1mM DTT, 0.2 U/ml Rnase inhibitor in 3X SSC buffer). The cell suspension was filtered through a 50-μm cell strainer. Processing for single-cell capture and library preparation was performed according to the 10X Genomics standard protocols.

### Immunostaining of mouse retinal sections

The dissection of mice retinas and eyecups was performed as previously described (Toulis et al., 2016). Cryosections of mouse retinas (10–12 μm slides) were obtained using a Leica CM3050-S cryostat. Immunostaining was performed as reported elsewhere (Aísa-Marín et al., 2020a). Antibodies were used following the subsequent dilutions: anti-GNAT2 (Abcam ab97501, 1:300), anti-RHO (Abcam ab5417, 1:300), anti-CRYBB2 (Santa Cruz sc-376006, 1:200), anti-S-OPSIN (Millipore AB5407, 1:300), anti-M-OPSIN (Millipore AB5405, 1:300), anti-CTSB (Abcam ab58802, 1:200), anti-MLKL 42hosphor S358 (Abcam ab187091, 1:200), anti-PAR (Abcam ab14459, 1:200). Slides were mounted using ProLong™ Gold Antifade Mountant (Thermo Fisher Scientific), and samples were kept refrigerated until confocal microscopy (Carl Zeiss LSM880).

### Electrophoresis of proteins in polyacrylamide gels (SDS-PAGE), Western blot and immunodetection

Retinas were lysed in radioimmunoprecipitation assay (RIPA), and electrophoresis was performed as reported (Aísa-Marín et al., 2020a). Following the manufacturer’s instructions, proteins were transferred to a PVDF Western Blotting Membrane (BioRad). The membrane was blocked with 10% non-fat dry milk in PBS containing 0.1% Tween 20 (Sigma-Aldrich). Primary antibodies (anti- GNAT2 [Abcam ab97501, 1:1000], anti-CRYAA [Abcam ab5595, 1:1000], anti-VDAC [Calbiochem Ab-5 185-197, 1:7000], rodent total OXPHOS cocktail [MitoSciences 6μg/ml]) were incubated overnight at 4°C followed by incubation with the corresponding secondary antibodies for 1 hour at room temperature using standard procedures. Proteins were visualized using the Luminescent Image Analyzer LAS-3000 mini (Fujifilm, Tokyo, Japan).

### RNA isolation and reverse transcriptase PCR (RT-PCR)

Wildtype mouse retinas were homogenized using a Polytron PT1200E homogenizer (Kinematica, AG, Lucerne, Switzerland). Total RNA was isolated using the Rneasy Mini Kit (Qiagen, Germantown, MD) and Rneasy Plus Mini Kit (Qiagen, Germantown, MD), following the manufacturer’s instructions with minor modifications (treatment with DNAse I during 1 h). Reverse transcription reactions were carried out using the qScriptTM cDNA Synthesis Kit (Quanta BioSciences, Inc., Gaithersburg, MD). Specific primers for amplification were designed and optimized. RT-PCR was performed according to standard thermocycling conditions.

### Primary retinal cultures

Primary retinal cultures were obtained from the dissection of P0-1 mouse retinas and processed as described in (Mirra et al., 2021). Briefly, the cell suspension was obtained using Neural Tissue Dissociation Kit (Miltenyi Biotec, Germany). 150,000 cells were plated onto poly-D-ornithine-laminin coated coverslips in Neurobasal™-A medium (Thermo Fisher Scientific) supplemented with 0.06% glucose, 0.0045% NaH2CO3,1 mM l-glutamine, B27 (Invitrogen), penicillin/streptomycin (Invitrogen 1%), forskolin 5 μM (Sigma Aldrich), BDNF 5 ng/ml (Peprotech) and rat CNTF 20 ng/ml (Peprotech). After 7 days of differentiation in vitro (7DIV), cells were fixed and stored in cryoprotection solution (30% Glycerol, 25% Etilenglicol 100% and 0.1 M PBS) for further experiments.

### Single-cell RNA-seq analysis

#### Pre-processing

Raw reads were trimmed using fastp (version 0.20.0) (Chen et al., 2018). 10x Genomics Cell Ranger (version 6.0.1) (Zheng et al., 2017) was used for barcode processing, alignment to the mouse reference genome (GENCODE vM23/Ensembl 98; https://support.10xgenomics.com/single-cell-gene-expression/software/downloads/latest?), and feature-barcode matrices generation. Cells were filtered by the number of counts and number of features to remove the top and bottom 5% of cells representing outliers (Luecken and Theis, 2019). Doublets were detected and filtered using DoubletFinder (version 2.0.3) (McGinnis et al., 2019).

Initial data examination revealed a broad presence of rod-cell marker expression throughout all individual cells, which was attributed to ambient RNA contamination from this abundant and morphologically challenging cell population. SoupX (version 1.5.2) (Young and Behjati, 2020) was used prior to cell filtering to remove ambient RNA using the functions `autoEstCont(sc, forceAccept=TRUE)` and `adjustCounts(sc)`. After cleaning ambient RNA, expression of specific gene markers (e.g. *Nrl* and *Rho* for rods) was highly enriched in the corresponding retinal cell population as shown in the UMAP plots (Fig. 1). Data is available at Suppl. Data 1.

#### Integration and clustering

The R package Seurat (version 4.1.0) (Butler et al., 2018; Stuart et al., 2019) was used to integrate our samples with a publicly available scRNA-seq dataset for mouse retina (Norrie et al., 2019). The use of a trusted reference to integrate our data ensured a better cell clustering identity and downstream analysis robustness of our scRNA-seq data. Sample integration was performed using the SCT normalization method. SCT transformed data were clustered using the first 30 principal components (PCs) and a resolution of 0.25.

#### Cell type identification and differential expression

RNA assay from the Seurat object was normalized and scaled to detect cell type markers. A set of variable features was identified using `selection.method=“vst”`. The `FindAllMarkers` function was then used to detect cluster-specific markers. Cluster cell identity was manually determined using known cell type markers (Hoang et al., 2020). Annotated Seurat clusters confidently sharing cell type markers were joined for downstream analysis. Misregulated genes for each cell type between wt samples and mutants were identified using the `FindMarkers` function. Resulting p-values were adjusted for multiple testing using `p.adjust(method=“fdr”)` as implemented in R (version 4.0.1).

#### Cone and rod sub-clustering

Cells identified as cone or rod were subset separately and re-clustered using SCT transformed data taking the first 20 PCs and a resolution of 0.01. Cell type sub- cluster markers were identified using RNA assay and `FindAllMarkers` as implemented in Seurat. Data is available in Suppl. Data 2 and Suppl. Data 3.

#### Expression visualization

Violin plots and UMAPs showing gene expression were produced using the counts of the Seurat SCT assay. Violin plots were log-transformed for visual clarity. UMAP expression plots were produced with the Seurat function `FeaturePlot` with the following arguments: `slot=“counts”`, `cols=c(“grey”,“red”)`, `keep.scale=NULL` or `“feature”`, `order=TRUE`, `max.cutoff=“q95”`.

#### RNA velocity

Unspliced and spliced mRNAs quantification was performed using velocyto (version 0.17.17)(La Manno et al., 2018) with the same reference genome annotation used for Cell Ranger and the mm10 repeat masker annotations downloaded from the UCSC table browser. RNA velocities were calculated using the scVelo (version 3.8.6)(Bergen et al., 2020) package in python (version 3.8.6) in the stochastic. To capture genotype specific RNA velocities, those were calculated for wt and mutant samples independently. RNA velocities for sub- clusters were calculated in the same way after the subsetting for rod or cone cells.

### Quantification and statistical analysis

Statistical significance of data, equal standard deviation (SD) and normal distribution were first assessed using Bartlett and Shapiro-Wilk tests. If data followed a normal distribution and showed homogeneity of variance, one-sample T-test was used for statistical significance analysis. When data did not follow a normal distribution, non-parametrical Mann-Whitney U-test was applied. Analysis was performed using GraphPad Prism 7.03 (San Diego, CA, USA).

## SUPPLEMENTAL INFORMATION TITLES AND LEGENDS

**Supplementary data 1. Filtered and processed data from the scRNA-seq analysis for the whole retina**. Included files are sparse matrix with RNA counts (after SoupX ambient RNA correction) in 10X format (matrix.mtx.gz, barcodes.tsv.gz, features.tsv.gz) that can be read with the `DropletUtils::read10xCounts` function. Metadata file with the information of cell type and genotype for each barcode.

**Supplementary data 2. Same as supplementary data 1 but for cone sub- cluster cells only.**

**Supplementary data 3. Same as supplementary data 1 but for rod sub- cluster cells only.**

**Table S1. Excel File with the list of DE genes grouped by their behaviour pattern and listed as presented in** **Figure 2A**. GO_IDs and GO_names columns contain the gene ontology IDs and names respectively of the gene. If more than one GO term is associated to the gene, then they are separated by a semicolon.

**Figure S1. Expression of specific marker genes enabled clear assignment of the main retinal cell types from scRNA-seq data.** Violinplots showing the main marker genes for the 7 cell types identified in the scRNA-seq data. The y-axis values correspond to log-transformed counts.

**Figure S2. Expression of *Nr2e3* in the wildtype and mutant retinas. A.** IGV visualization of scRNA-seq reads pileup in the cone and rod cell populations aligned to the reference mouse genome validates the gene-editing alteration in the genotype of the *Nr2e3* mutants. The deletion of 27 nucleotides in exon 8 in the Δ27 mutant is detected by a sharp blank in the RNA reads alignment (indicated by a black arrowhead). In contrast, the ΔE8 mutant shows complete deletion of exon 8 (no reads are aligned) and different transcript reads with partial intron 7 retention. The scale of the number of reads in rods and cones is different and the results should be interpreted only qualitatively. **B.** Quantification of the expression of *Nr2e3* in the rod and cone subclusters per each biological sample (3 wt retinas, and 2 of each mutant genotype). Δ27 *Nr2e3* mutants show a 3.5- fold increase in *Nr2e3* expression in both rods and cones compared to the wt. In the ΔE8 mutant, the expression of *Nr2e3* is shallow in the two photoreceptor types. **C.** Ratios of *Nr2e3* expression in cone and rod photoreceptors (per each biological sample) show that approximately 75% of the total expression is due to rods, and 25% to cones, which is maintained in the three genotypes. **D.** The long isoform protein is more stable than that encoded in the short isoform. Protein quantification in cells transfected with either the *Nr2e3* long or short isoforms (treated or untreated with 10 mM MG132 for 16 h) shows that the stability of the short isoform is reduced compared to the long isoform. Total cell lysates were subjected to western blot analysis with anti-NR2E3 and quantified. Values were plotted after normalization with β-tubulin and represented the fold change compared to the long isoform values in cells untreated with the MG132 proteasome inhibitor (–MG132). Data represent the mean ± SD of 5 independent experiments (*P<0.05).

**Figure S3. Differential expression of retinal specific genes related to light perception, between mutants and wt in cone and rod cells.** Log2-fold change heatmap of DE genes related to photoreceptor function based on their GO tems.

**Figure S4. Expression of rod and cone genes in global rod and cone populations separated by genotype.** Note the higher expression of most rod genes in the rods of the mutant vs wt retinas, but not as much in the cones (*Gnat1, Gngb1, Gngt1, Rho, Rom1, Pdc*). On the other hand, some of the cone-specific genes are also more highly expressed in the mutant than wt cones (*Gnat2, Gnb3, Opn1sw, Pde6c, Pde6h*). Some of the cone genes are also highly dysregulated and expressed in the rods of the mutant retinas (*Gnat2, Gnb3, Pde6c, Pde6h*). The highest expression of *Nr2e3* in both rods and cones of the D27 mutant deserves specific mention. The y-axis values correspond to log-transformed counts.

**Figure S5. Overexpression of foveal marker genes is increased in the mutant retinas.** Overexpression of the foveal marker genes *Gngt1* and *Cyp26a1* in the mutants was detected by RT-PCR, thus confirming some of the genes highlighted after the analysis of scRNA-Seq data. Samples 1 to 10 correspond to independent biological replicates from the three genotypes (1-3 wt, 4-7 Δ27, 8- 10 ΔE8). Molecular weight marker used in A and B is a 100 bp-ladder (NZYDNA Ladder V, Nzytech).

**Figure S6. Expression of cone, rod, and photoreceptor differentiation genes in the cone subclusters by genotype.** Cone genes are highly expressed in subcluster cone_0_, whereas rod genes are more expressed in subclusters cone_1_ and cone_4_. Subclusters cone_2_ and cone_3_ show higher expression of genes related to photoreceptor differentiation. The y-axis values correspond to log-transformed counts.

**Figure S7. Gene expression and GO term analysis of rod subclusters. A.** Log2-fold change heatmap of rod subcluster marker genes sorted by k-means clustering. Frequent biological process GO terms appear in the cloud according to their presence in each cluster. **B.** Expression of rod- (*Gngt1, Gnat1, Rho, Nr2e3*) and cone-specific genes (*Gnb3, Pde6h, Opn1sw, Gnat2*) in each rod subcluster by genotype. *Nr2e3* is overexpressed in the Δ27 rods compared to the wt and the ΔE8 rods. Cone genes are significantly upregulated in almost all rod subclusters in the mutant retinas. The y-axis values correspond to log-transformed counts.

**Figure S8. UMAP plot showing the differential expression of S- and M-opsin.** The population of cones (arrows in the magnifications) that show directional flow towards the rod cluster (main figure 4) correspond to S-cones, as inferred from the expression of S-opsin (Opn1sw) but not M-opsin (Opn1mw).

## Notes

### Competing Interest Statement

The authors have declared no competing interest.

