## Supplemental Figures 1-8 for "Specific photoreceptor cell fate pathways are differentially altered in NR2E3-associated diseases"

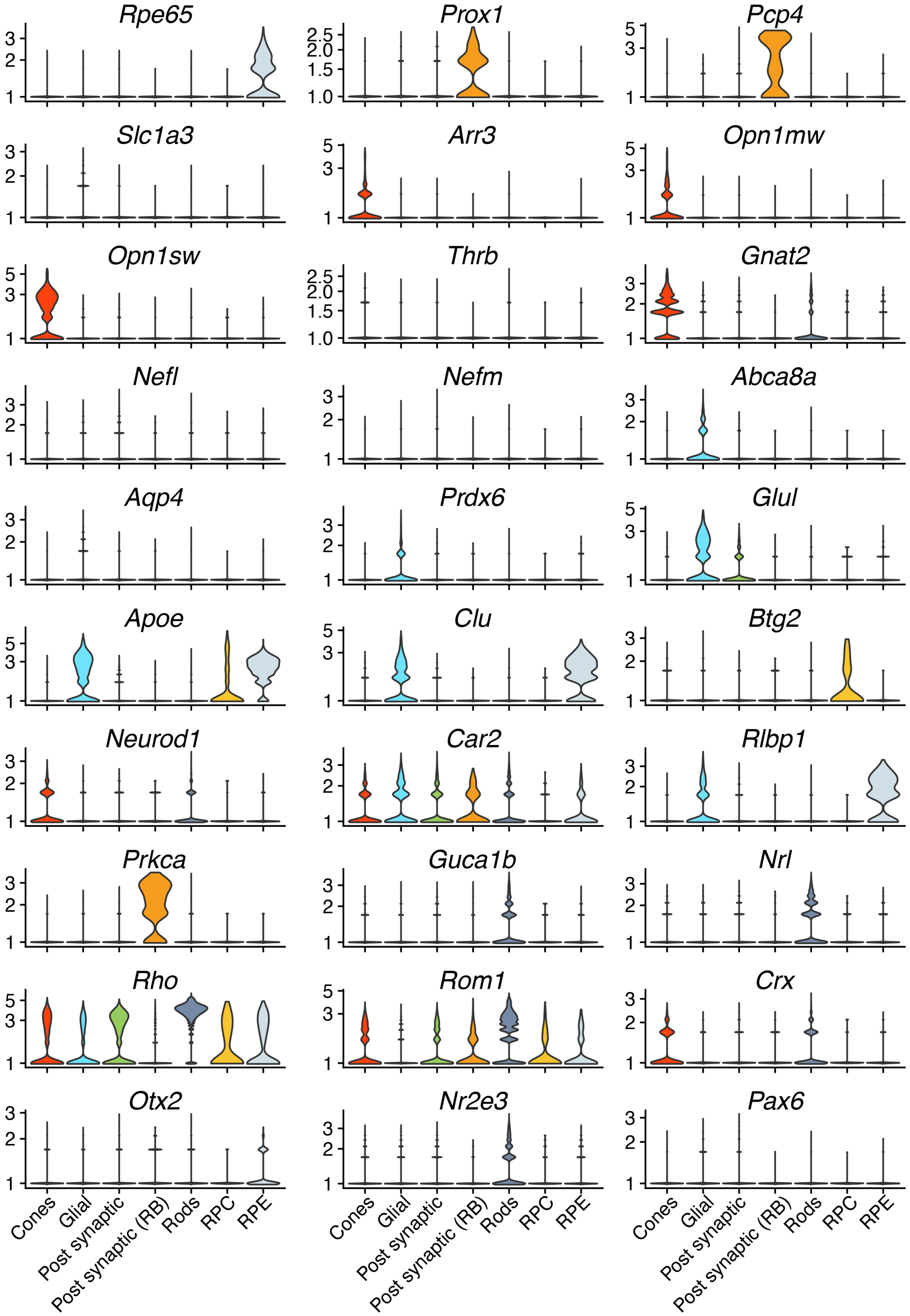


**Figure S1. Expression of specific marker genes enabled clear assignment of the main retinal cell types from scRNA-seq data.** Violin plots showing the main marker genes for the 7 cell types identified in the scRNA-seq data. The y-axis values correspond to log-transformed counts.

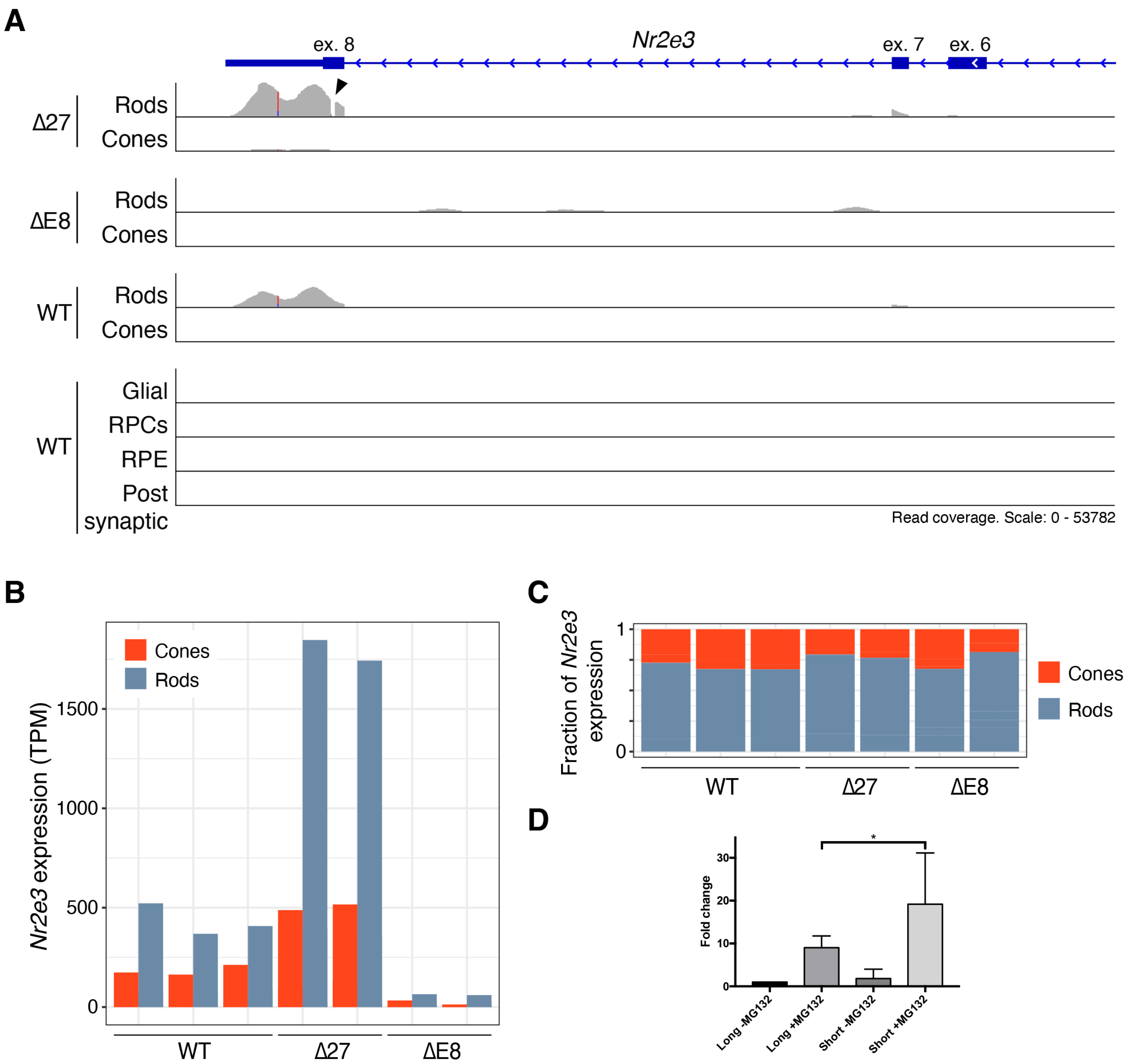


**Figure S2. Expression of *Nr2e3* in the wildtype and mutant retinas. A.** IGV visualization of scRNA-seq reads pileup in the cone and rod cell populations aligned to the reference mouse genome validates the gene-editing alteration in the genotype of the *Nr2e3* mutants. The deletion of 27 nucleotides in exon 8 in the Δ27 mutant is detected by a sharp blank in the RNA reads alignment (indicated by a black arrowhead). In contrast, the ΔE8 mutant shows complete deletion of exon 8 (no reads are aligned) and different transcript reads with partial intron 7 retention. The scale of the number of reads in rods and cones is different and the results should be interpreted only qualitatively. **B.** Quantification of the expression of *Nr2e3* in the rod and cone subclusters per each biological sample (3 wt retinas, and 2 of each mutant genotype). Δ27 *Nr2e3* mutants show a 3.5-fold increase in *Nr2e3* expression in both rods and cones compared to the wt. In the ΔE8 mutant, the expression of *Nr2e3* is shallow in the two photoreceptor types. **C.** Ratios of *Nr2e3* expression in cone and rod photoreceptors (per each biological sample) show that approximately 75% of the total expression is due to rods, and 25% to cones, which is maintained in the three genotypes. **D.** The long isoform protein is more stable than that encoded in the short isoform. Protein quantification in cells transfected with either the *Nr2e3* long or short isoforms (treated or untreated with 10 mM MG132 for 16 h) shows that the stability of the short isoform is reduced compared to the long isoform. Total cell lysates were subjected to western blot analysis with anti-NR2E3 and quantified. Values were plotted after normalization with β-tubulin and represented the fold change compared to the long isoform values in cells untreated with the MG132 proteasome inhibitor (–MG132). Data represent the mean ± SD of 5 independent experiments (*P<0.05).

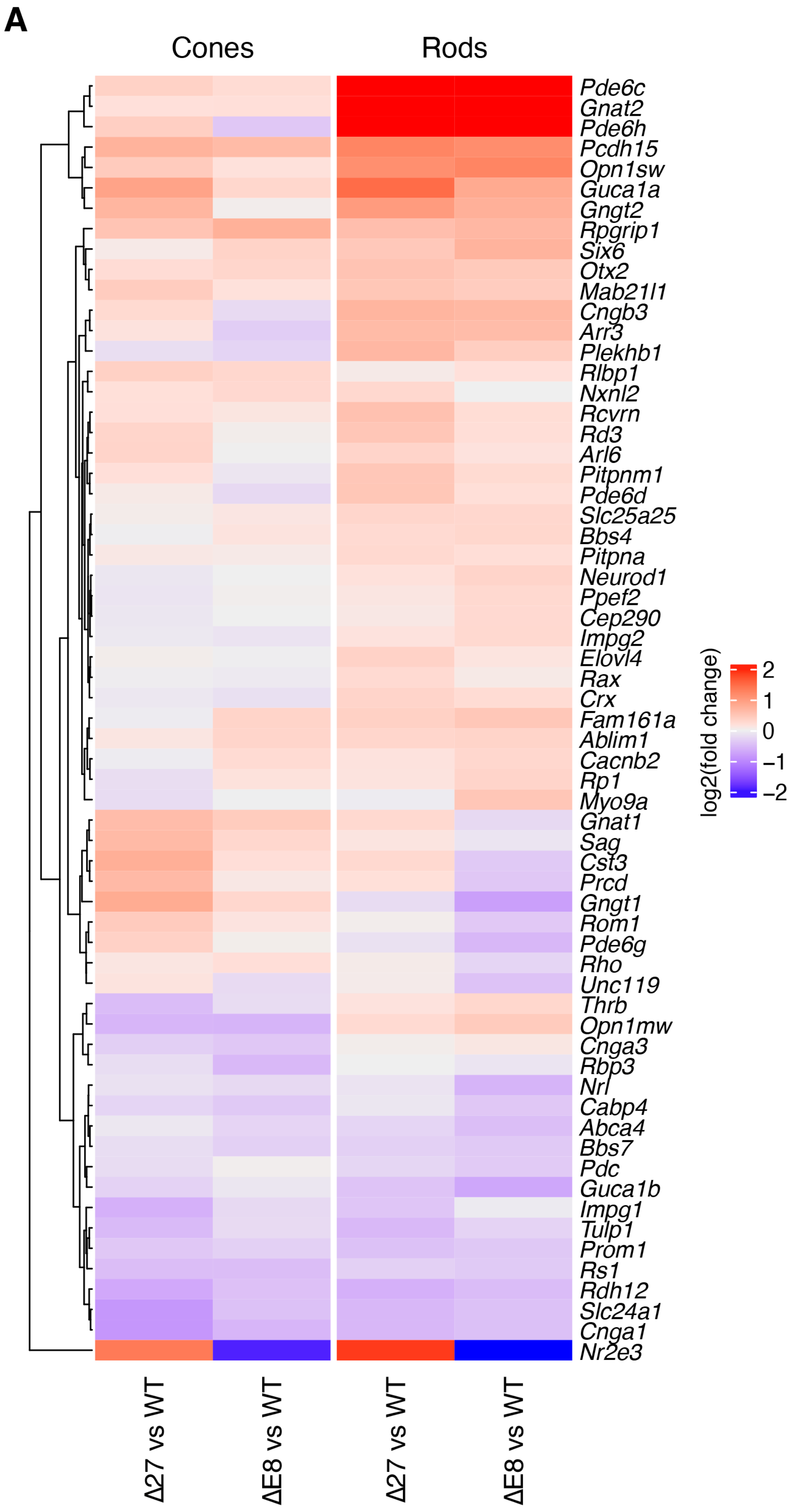


**Figure S3. Differential expression of retinal specific genes related to light perception, between mutants and wt in cone and rod cells.** Log2-fold change heatmap of DE genes related to photoreceptor function based on their GO tems.

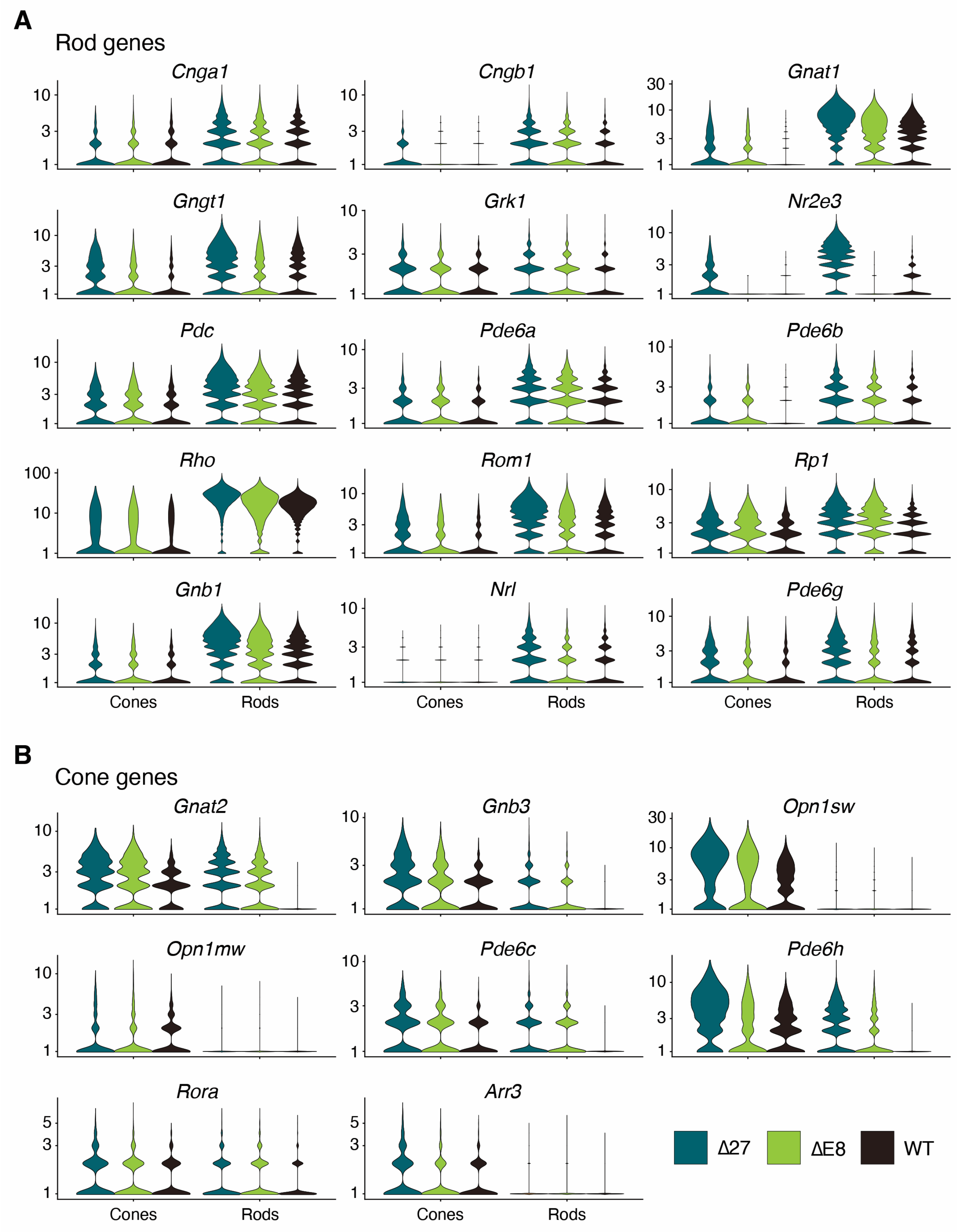


**Figure S4. Expression of rod and cone genes in global rod and cone populations separated by genotype.** Note the higher expression of most rod genes in the rods of the mutant vs wt retinas, but not as much in the cones (*Gnat1, Gngb1, Gngt1, Rho, Rom1, Pdc*). On the other hand, some of the cone-specific genes are also more highly expressed in the mutant than wt cones (*Gnat2, Gnb3, Opn1sw, Pde6c, Pde6h*). Some of the cone genes are also highly dysregulated and expressed in the rods of the mutant retinas (*Gnat2, Gnb3, Pde6c, Pde6h*). The highest expression of *Nr2e3* in both rods and cones of the D27 mutant deserves specific mention. The y-axis values correspond to log-transformed counts.

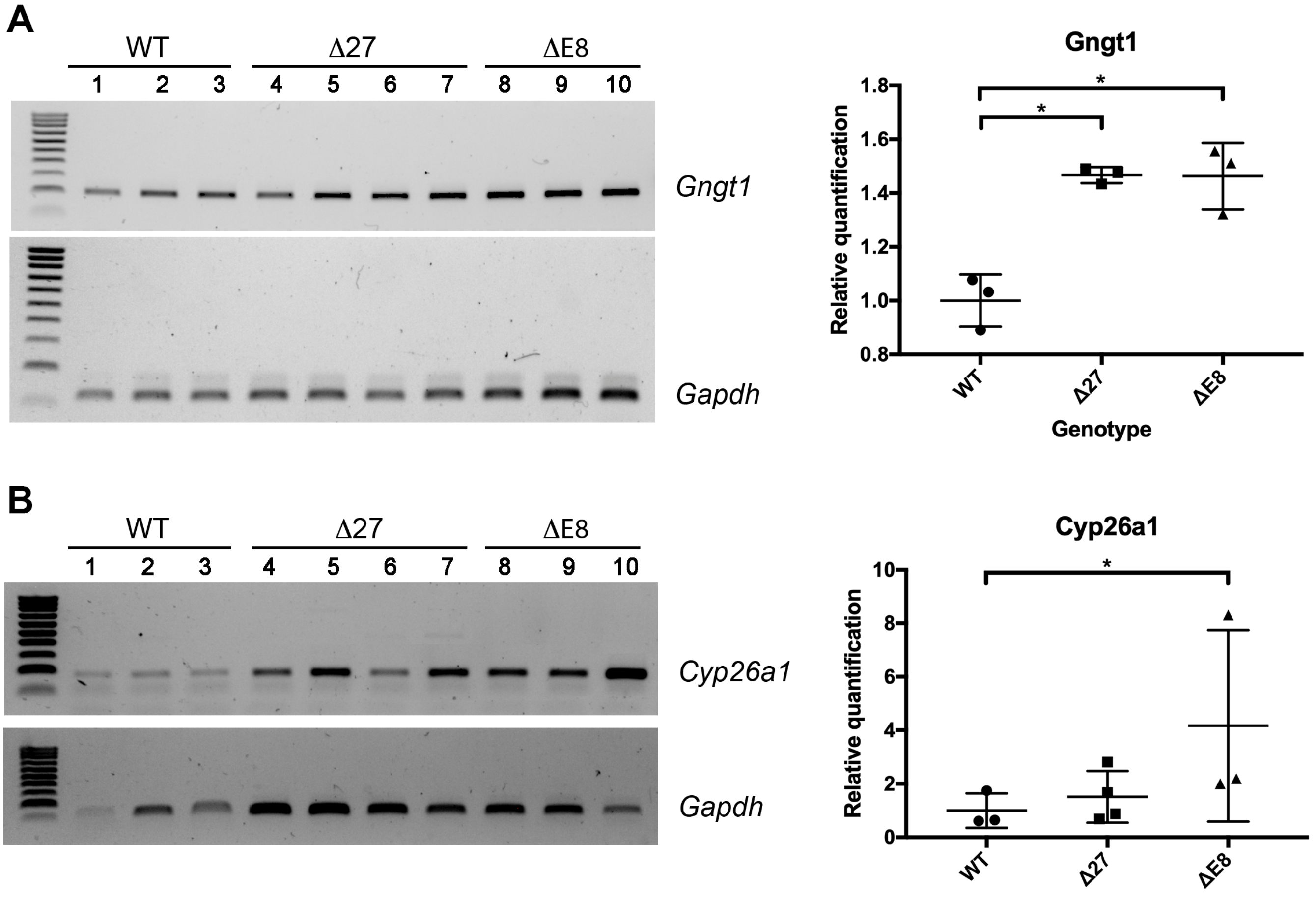


**Figure S5. Overexpression of foveal marker genes is increased in the mutant retinas.** Overexpression of the foveal marker genes *Gngt1* and *Cyp26a1* in the mutants was detected by RT-PCR, thus confirming some of the genes highlighted after the analysis of scRNA-Seq data. Samples 1 to 10 correspond to independent biological replicates from the three genotypes (1-3 wt, 4-7 Δ27, 8-10 ΔE8). Molecular weight marker used in A and B is a 100 bp-ladder (NZYDNA Ladder V, Nzytech).

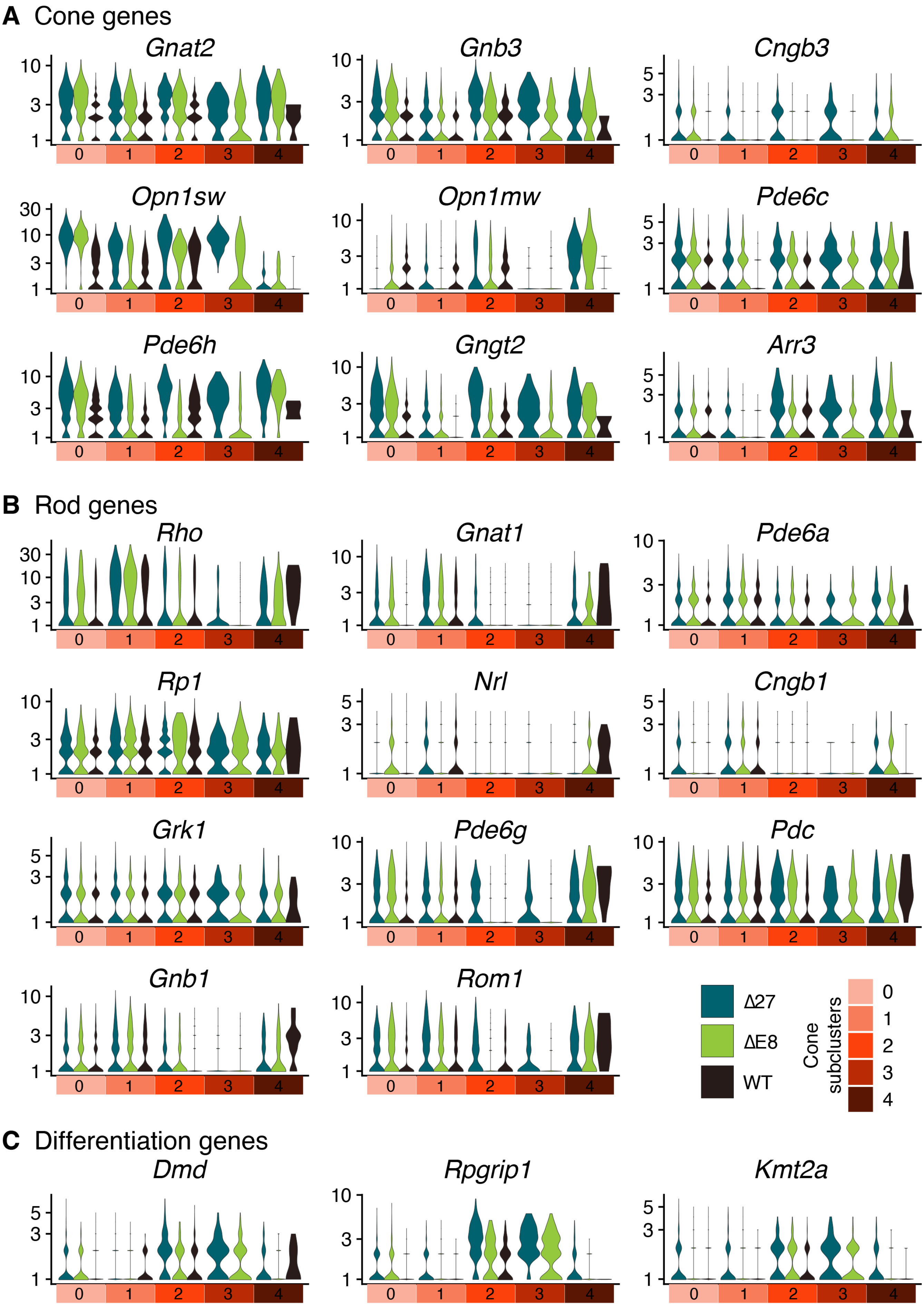

A Cone genes
Gnat2
Gnb3
Cngb3
Opn1sw
Opn1mw
Pde6c
Pde6h
Gngt2
Arr3
B Rod genes
Rho
Gnat1
Pde6a
Rp1
Nrl
Cngb1
Grk1
Pde6g
Pdc
Gnb1
Rom1
C Differentiation genes
Dmd
Rpgrip1
Kmt2a
Δ27
ΔE8
WT
Cone subclusters
0
1
2
3
4

**Figure S6. Expression of cone, rod, and photoreceptor differentiation genes in the cone subclusters by genotype.** Cone genes are highly expressed in subcluster $cone_0$, whereas rod genes are more expressed in subclusters $cone_1$ and $cone_4$. Subclusters $cone_2$ and $cone_3$ show higher expression of genes related to photoreceptor differentiation. The y-axis values correspond to log-transformed counts.

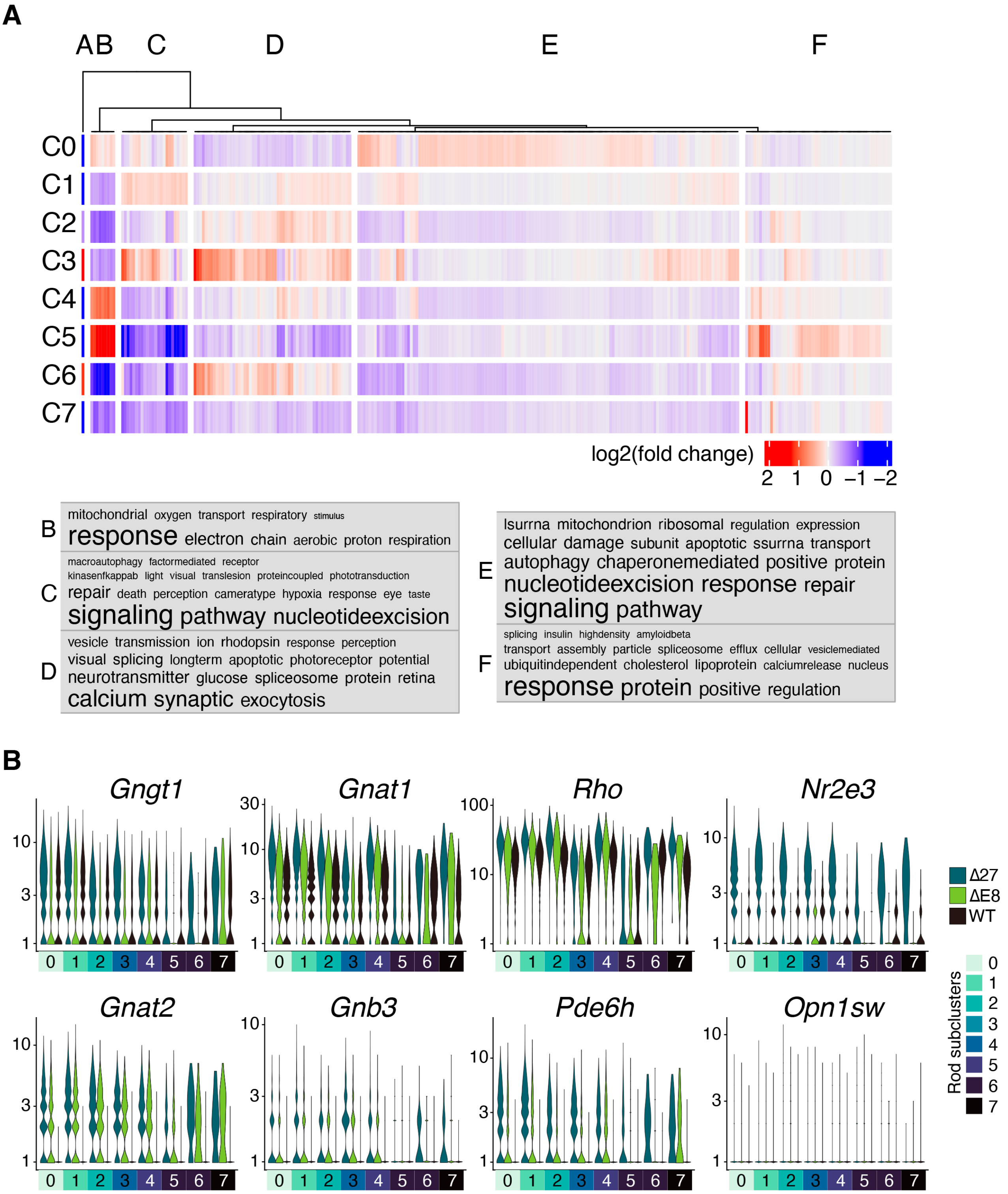


**Figure S7. Gene expression and GO term analysis of rod subclusters. A.** Log2-fold change heatmap of rod subcluster marker genes sorted by k-means clustering. Frequent biological process GO terms appear in the cloud according to their presence in each cluster. **B.** Expression of rod- (*Gngt1, Gnat1, Rho, Nr2e3*) and cone-specific genes (*Gnb3, Pde6h, Opn1sw, Gnat2*) in each rod subcluster by genotype. *Nr2e3* is overexpressed in the Δ27 rods compared to the wt and the ΔE8 rods. Cone genes are significantly upregulated in almost all rod subclusters in the mutant retinas. The y-axis values correspond to log-transformed counts.

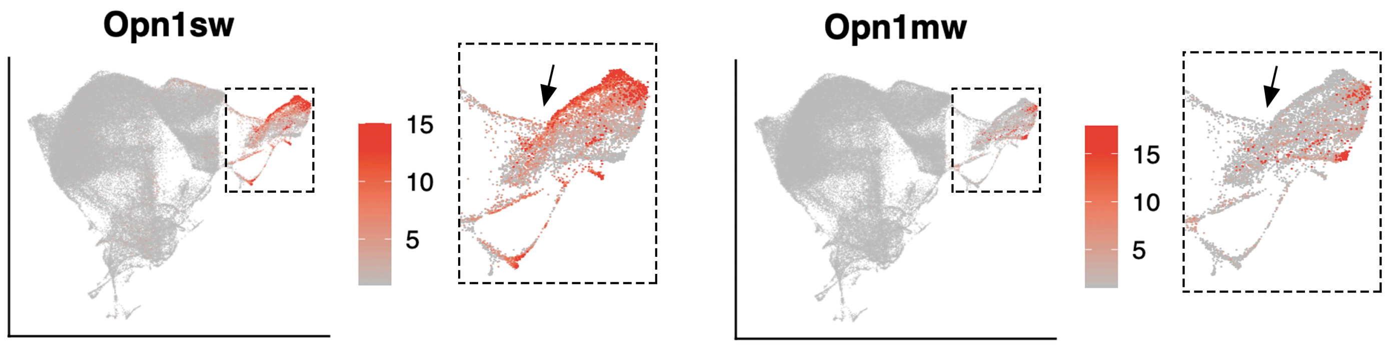


**Figure S8. UMAP plot showing the differential expression of S- and M-opsin.** The population of cones (arrows in the magnifications) that show directional flow towards the rod cluster (main figure 4) correspond to S-cones, as inferred from the expression of S-opsin (Opn1sw) but not M-opsin (Opn1mw).
